# CD14 plays a critical role in pain and inflammation across multiple models of post-traumatic osteoarthritis

**DOI:** 10.1101/2025.06.02.657478

**Authors:** Kevin G. Burt, Sanique M. South, Natalie S. Adamczyk, Vu Nguyen, Shingo Ishihara, Cheng Zhou, Kate L. Sharp, Sung Y. Kim, Anna E. Rapp, Baofeng Hu, Lance A. Murphy, Li-Yin Hung, De’Broski Herbert, Joshua F. Baker, Robert L. Mauck, Timothy M. Griffin, Anne-Marie Malfait, Rachel E. Miller, Carla R. Scanzello

## Abstract

Inflammation is a primary driver of osteoarthritis (OA), and no therapies exist to halt or delay disease progression or substantially ameliorate the chronic pain, inflammation and disability that are characteristic of disease. Soluble CD14 (sCD14), a co-factor that enhances inflammatory toll-like receptor signaling, is present in synovial fluid in patients with OA and positively associates with joint space narrowing, synovial macrophage content, and pain. In this study, we show that increased sCD14 within human synovial fluid correlates with joint effusion volume and increased knee hyperalgesia. Next, we evaluated CD14 as a potential therapeutic target in three pre-clinical models of post traumatic OA (PTOA), using both genetic deficiency and pharmacologic blockade to modulate its activity. We demonstrate that deficiency or blockade of CD14 results in significant protection from increased evoked pain behaviors and from OA driven mobility impairments (i.e. decreased cage activity) across models that differ in severity, and across male and female cohorts. Using flow cytometry, single cell transcriptomics, and spatial proteomics, we further show that CD14-deficiency drastically influences the local synovial inflammatory landscape post-injury, reducing monocyte and macrophage populations and modulating local fibroblast populations. Targeting CD14 via genetic deficiency or therapeutic blockade revealed no substantial protection, but no worsening, of cartilage degeneration. Ultimately, our results provide strong support that targeting synovial inflammation through blockade of CD14 can safely ameliorate OA pain and disability after a pre-disposing injury.

**One Sentence Summary:** The study demonstrates the key role of CD14 in pain, mobility loss, and inflammation in PTOA, and demonstrates the therapeutic potential of CD14 blockade for OA pain relief.

## INTRODUCTION

Osteoarthritis (OA) is a degenerative joint disease characterized by joint pain and loss of mobility, ultimately causing a substantial disruption to daily life in millions of people each year (*1*). Knee OA is characterized pathologically by the breakdown of articular cartilage and accompanying synovial inflammation. The synovium, which normally produces factors that lubricate the joint to facilitate proper knee function, becomes infiltrated by immune cells during OA progression, with the most prevalent being monocyte/macrophages (*2*). Though non-steroidal anti-inflammatory medications can provide temporary pain relief, these drugs have significant side effects and do not specifically target disease mechanisms. No therapies have been able to halt or delay disease progression or prevent the onset of chronic pain and disability (*3*). Given that inflammation is a central feature of OA that is strongly associated with pain and joint dysfunction, as well as progression of cartilage loss, many pre-clinical studies are beginning to evaluate targeting synovial inflammation specifically for the treatment of painful OA (*3*).

Previously, our group identified a possible mediator of synovial inflammation in OA, soluble CD14 (sCD14) (*4*). Others subsequently showed that synovial fluid levels of sCD14 are positively associated with joint space narrowing and pain (*5*). sCD14 is released from monocytes following inflammatory activation and correlates with synovial macrophage content in OA patients (*5–7*). sCD14 is thought to act as a damage associated molecular pattern (DAMP), interacting with the pattern recognition toll like receptor 4 (TLR4) to initiate broad inflammatory pathway activation via NF-κB (*8*). While surface membrane bound CD14 (mCD14) is most highly expressed on monocyte/macrophage populations, it can also be shed by these cells to transactivate a host of additional immune cells (e.g. neutrophils, dendritic cells, T-cells) and non-immune cell types (e.g. epithelial, endothelial cells) (*9*). On myeloid cells, mCD14 functions to promote binding of lipopolysaccharide (LPS), serving as a co-receptor and facilitating recognition by TLR4 and subsequent downstream transcription of a broad array of inflammatory cytokines (*9*). In addition, CD14 can enhance inflammatory signaling in response to other TLR ligands, including pathogen associated molecular pattern (PAMPs) like lipoteichoic acid, and damage associated molecular patterns (DAMPs) including S100 calcium binding protein A9 (S100A9), biglycan, and high mobility group box 1 (HMGB1) (*10*).

We previously reported that global genetic CD14-deficiency in mice protects against OA-associated bone-remodeling and cartilage damage and mitigated mobility loss during OA progression in a model of post-traumatic OA (*11*). These results provided a strong rationale for investigating how these structural and functional protections relate to pain behaviors and whether CD14 can be targeted therapeutically. Specifically, we aimed to test the hypothesis that CD14 inhibition attenuates synovial inflammation and associated pain during disease progression.

Multiple pre-clinical models have been established that reproduce OA disease pathology and progression (e.g. inflammation, cartilage loss, pain). Though reproducible, the use of a single model is limited in recapitulating the complexity of OA commonly observed within the clinical population. Moreover, many of these models exhibit sexually dimorphic disease pathology and symptoms. Towards translation, we employed both global genetic deletion of CD14 and intra-articular CD14 blockade across multiple murine OA models that vary in severity of pathology and rate of progression. We first demonstrated that soluble CD14 positively correlates with increased joint inflammation and pain in OA patients. Next, we showed that a global CD14-deficiency mitigates spontaneous and evoked pain responses across multiple murine models of OA, reduces synovial monocyte populations, and modulates synovial fibroblast populations during OA. Lastly, we demonstrated the therapeutic potential of a CD14 neutralizing antibody, delivered locally to the joint shortly following injury, in mitigating spontaneous and evoked pain responses across surgical and non-surgical models of murine PTOA.

## RESULTS

### Soluble CD14 (sCD14) correlates with increased joint effusion and pain in OA patients

Synovial fluid (SF) was obtained from a cohort of patients enrolled in the Marching On for VEterans with Osteoarthritis of the Knee (MOVE-OK) clinical trial (clinicaltrials.gov identifier NCT05035810) with painful knee OA who underwent knee joint aspiration prior to- and 16-wks following treatment with either lidocaine only, or lidocaine plus a corticosteroid (**Table S1**) (*12*). sCD14 protein levels (total ng) were quantified within SF by ELISA. Knee algometer measurements were taken prior to joint aspiration, and pain pressure thresholds were recorded (kg/cm^2^), with lower thresholds indicating greater knee hyperalgesia, or higher levels of pain sensitization. Independent of treatment (which remains blinded) sCD14 levels strongly correlated (Spearman r (r_s_) = 0.91, p<0.0001) with total joint effusion volume (mL), an indicator of joint inflammation (**Fig. 1A**) (*13*). Higher sCD14 levels also trended with lower knee algometer recorded pressure pain threshold values (increased pain) (Spearman r = -0.33, p=0.082) (**Fig. 1B**). Linear regression analysis was then applied to control for age, sex, and multiple samples from a single patient (**Table S1**) (i.e. 0-wk pretreatment and 16-wk post treatment). This analysis revealed sCD14 to be significantly correlated (p<0.0001) with SF effusion volume. For each log total ng sCD14 increase, the SF effusion volume was 11.2 mL greater (**Fig. 1A**). Analysis also revealed that sCD14 significantly correlated with pain sensitization (p=0.036) with a coefficient of -2.34, representing that per log total ng sCD14 increase, the algometer pain pressure threshold was 2.34 kg/cm^2^ lower (**Fig. 1B**).

**Fig. 1:**
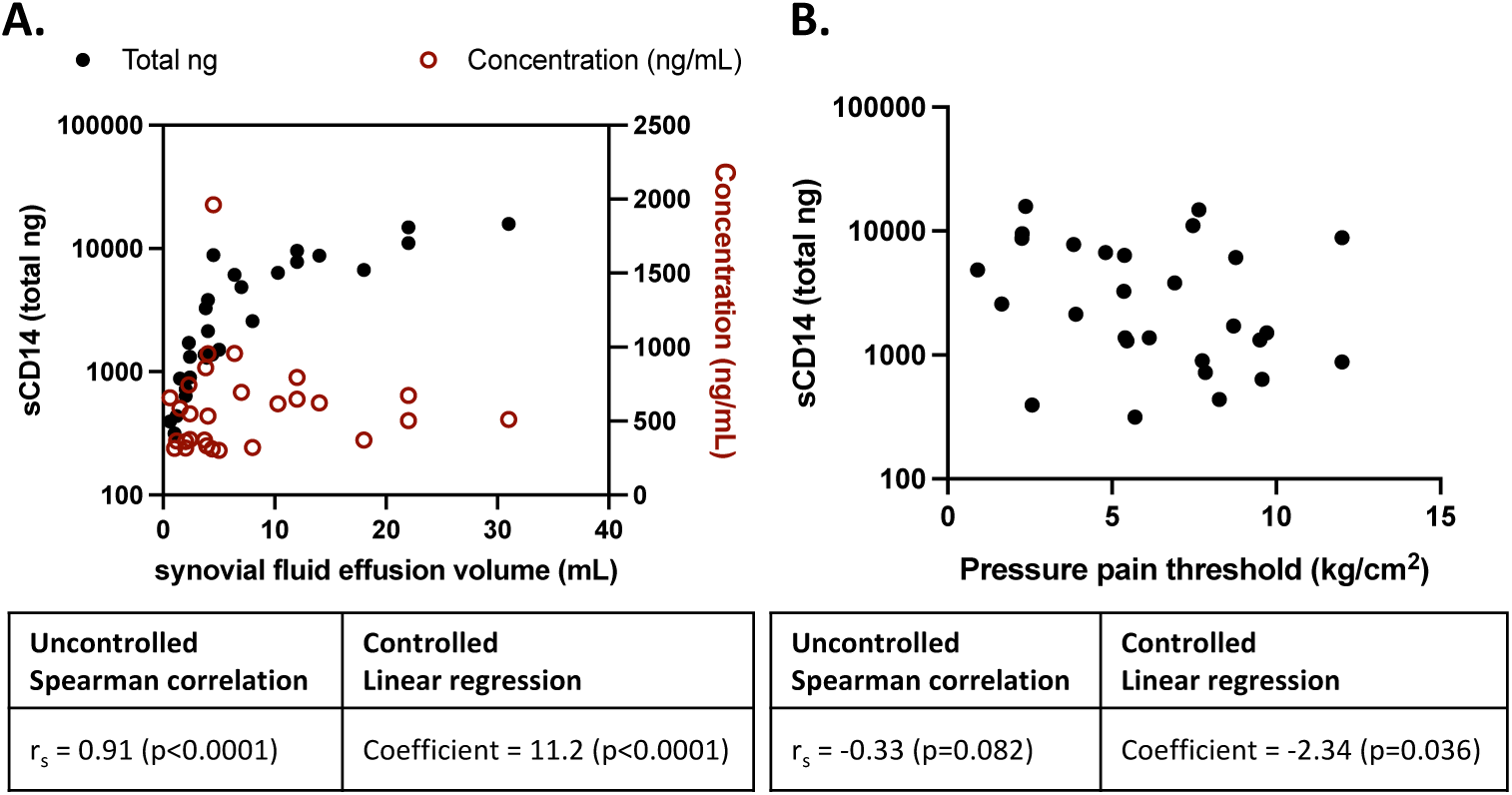
Evaluation of sCD14 expression within inflamed and painful knee joints. (**A**) SF fluid effusion volume (mL) plotted against total ng (black dots) and concentration (ng/mL, red circles) of sCD14 (n=14 patients, n=28 SF samples). (**B**) SF fluid effusion volume (mL) plotted against pressure pain threshold (kg/cm^2^). Spearman ranked correlation and controlled linear regression analysis of sCD14 levels (total ng) with total synovial fluid effusion volume (mL) and with knee algometer recorded pressure pain threshold values (kg/cm^2^). Spearman correlation (r_s_), linear regression coefficient, and significance (p) reported.

### Global CD14-deficiency mitigates spontaneous and evoked pain behaviors in two PTOA models

We next explored the role of CD14 in influencing synovial inflammation and symptomatic knee pain in preclinical models. Destabilization of the medial meniscus (DMM) surgery was performed in 12-wk old CD14 knockout (CD14 KO) and congenic wild-type (WT) male mice (*11, 14*). Evoked pain behavior (knee hyperalgesia) was measured by recording withdrawal forces (up to 450g) at the knee joint using a pressure application measurement (PAM, Ugo Basile) device. DMM surgery increased hyperalgesia (a decrease in withdrawal threshold) in WT mice at both 4-(p=0.0002) and 8-wks (p=0.01) post-injury, compared to pre-operative levels (**Fig. 2A**). In CD14 KO mice, DMM produced no significant knee hyperalgesia post-DMM (**Fig. 2A**) and, compared to WT mice, CD14 KO mice at 4-wks post-injury had significantly less hyperalgesia (p=0.00024, **Fig. 2A**). Next, decreases in rearing behavior and paw weight bearing asymmetry were measured using the advanced dynamic weight bearing (ADWB, Bioseb Inc.) system to evaluate spontaneous pain behavior. Paw weight bearing dynamics were assessed across a 5-minute testing period. CD14 KO mice exhibited greater rearing activity at pre-operative (p=0.0123) and 4-wks (p=0.04) post-DMM surgery timepoints compared to WT mice (**Fig. 2A**). DMM produced no significant changes to paw weight bearing within strain, and no changes to paw weight bearing was observed between CD14 KO and WT mice, including front to rear paw weight bearing ratios and rear right (injured) to rear left paw (uninjured) weight bearing ratios (**Fig. 2A**).

**Fig. 2:**
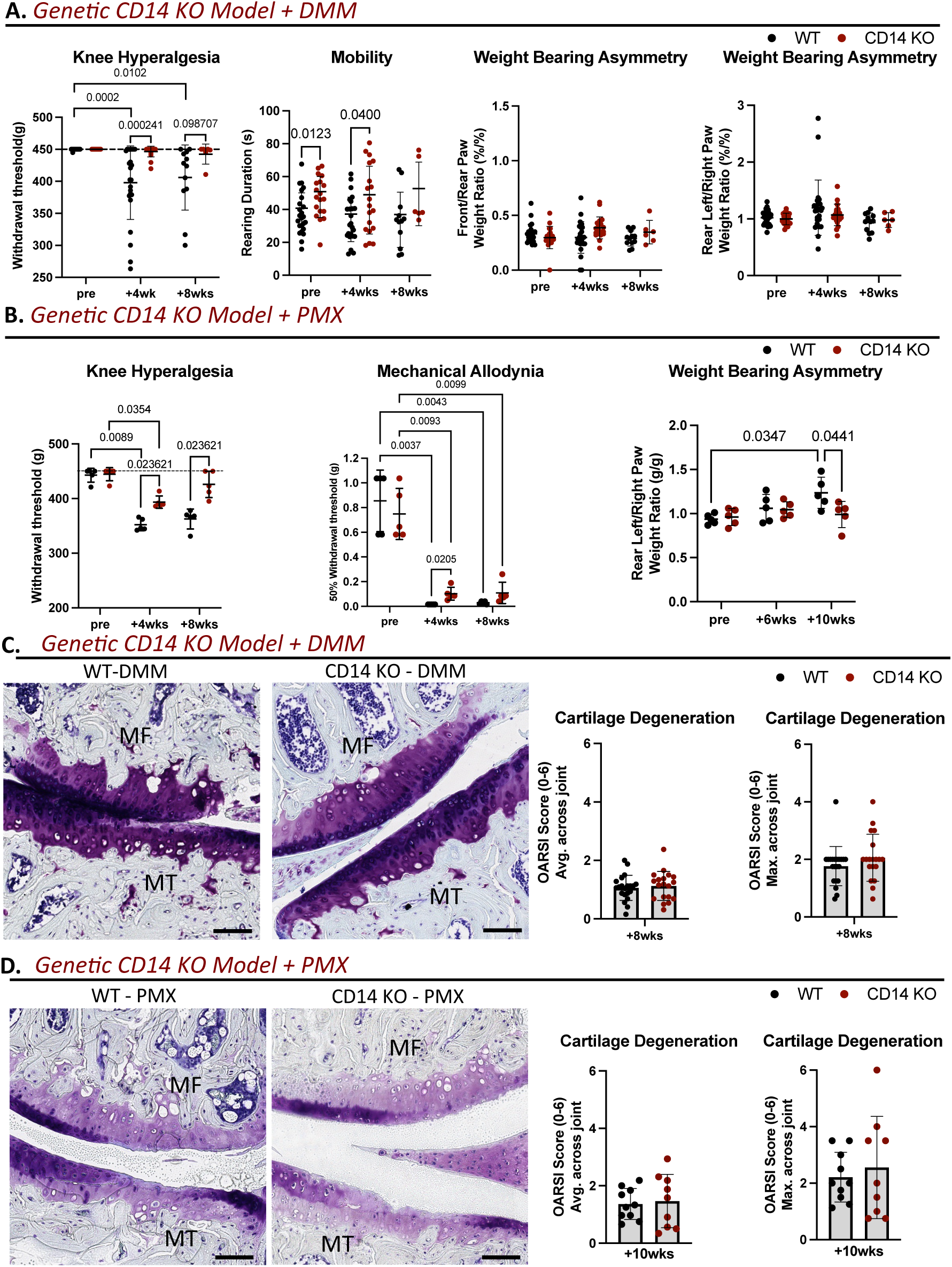
Pain behavior and cartilage damage following injury. (**A**) Pain, mobility, and weight bearing changes between WT (n=12-24) and CD14 KO (n=6-19) mice. Knee hyperalgesia measured via withdrawal threshold (g) using a PAM device. Mobility measured via rearing duration (s) using the ADWB system. Weight bearing asymmetry measured as the front to rear paw weight ratio (%/%) and rear-left to rear-right paw weight ratio (%/%). (**B**) Pain and weight bearing changes between WT (n=5) and CD14 KO (n=5) mice following PMX surgery. Knee hyperalgesia (evoked pain) measured via withdrawal threshold (g) using a PAM device. Mechanical allodynia measured via the 50% withdrawal threshold (g) using Von Frey. Weight bearing asymmetry measured as the rear-left to rear-right paw weight ratio (%/%) using the static incapacitance meter system. (**C, D**) Toluidine Blue stained histological sections of WT and CD14 KO tibial and femoral cartilage surfaces 8-wks following DMM (n=14-17) or 10-wks following PMX (n=9-10) (MF: medial femoral, MT: medial tibial) (Scale bar = 100μm). Statistically significant differences in knee hyperalgesia between groups (**A,B**) were calculated using the Mann-Whitney U non-parametric T-test with Holm-Šídák post-hoc, and across time points using either the Friedman test with Dunn’s multiple comparison adjustment or the Kruskal-Wallis test with Dunn’s multiple comparison adjustment for data sets with missing values. Statistically significant differences in remaining behavioral outcomes (**A,B**: mobility, allodynia, & weight bearing asymmetry) were calculated using a repeated measures two-way ANOVA or repeated measures mixed effect model followed by Šídák post-hoc for multiple comparisons. Statistically significant differences in histopathological scoring (**C,D**) were calculated using a student’s T-test.

Similar evoked and spontaneous pain behavior changes were next measured in a more severe surgical PTOA model generated via partial meniscectomy (PMX) in adult male mice of both strains (*15*). As expected, PMX produced a more severe pain phenotype than DMM, with a greater magnitude of hyperalgesia at the 4-wk post-injury time point in both CD14 KO (p=0.035) and WT (p=0.0089) mice compared to their respective pre-operative levels (**Fig. 2B**). Similar to results in the DMM model, CD14 KO mice developed significantly less hyperalgesia compared to WT at 4-(p=0.024) and 8-wks (p=0.024) post-PMX (**Fig. 2B**). Another evoked pain behavior, mechanical allodynia, was measured using von Frey fibers, recording withdrawal thresholds in response to microfilament probing of the rear hind paw on the PMX operated side (*16, 17*). Both CD14 KO and WT mice exhibited increased mechanical allodynia at 4- and 8-wks post-surgery (**Fig. 2B**). However, at the 4-wk time point, the 50% withdrawal threshold was significantly lower (indicating greater sensitivity) in WT compared to CD14 KO mice (p=0.02) (**Fig. 2B**). Lastly, PMX surgery produced a shift in rear paw weight bearing asymmetry in WT mice, with an increase on the left (unoperated) and a decrease on the right (PMX operated) leading to a change in the rear paw weight bearing ratio (p=0.035) at 10-wks compared to preoperative measurements (**Fig. 2B**). This rear left to right paw weight bearing ratio was significantly increased in WT mice compared to CD14 KO mice at 10 weeks post-PMX (p=0.044) (**Fig. 2B**). Notably, CD14 KO mice showed no change in rear left paw to rear right paw weight bearing ratio, or asymmetry, following PMX (**Fig. 2B**). The observed mitigations of pain behaviors and mobility within CD14 KO mice across models of ranging OA severity greatly supports a possible primary role of CD14 in painful OA progression.

We extended our evaluation in the PMX model to include a skeletally mature female cohort, given that both sexes develop robust structural disease pathology in this model (in contrast to DMM, where males develop OA symptoms more quickly) (33). Similar to our observations in male mice, CD14 KO female mice developed less knee hyperalgesia after PMX surgery with higher withdrawal thresholds observed at 4-(p=0.026) and 8-wks (p=0.034) compared to female WT mice (**Fig. S1**). Though nonsignificant, we observed a similar trend (p=0.072) of decreased weight bearing asymmetry within CD14 KO female PMX mice compared to female WT (**Fig. S1**). Adding to the clinical translatability, the observed pain behavior mitigations suggest a pivotal role of CD14 in PTOA associated pain across sexes.

### Global CD14 KO provided no protection to early-stage cartilage degeneration

Cartilage degeneration was evaluated on toluidine blue stained coronal knee sections taken midway through the joint using the modified Osteoarthritis Research Society International (OARSI) score (*18*). Histopathologic scoring was performed at early and mid-stage disease progression (4- to 8-wks) in the DMM model and at 8-wks post-surgery in the PMX model. No significant difference in cartilage degeneration scores were observed in CD14 KO mice at either 4- or 8-wk time points post-DMM compared to the WT cohort (**Fig. S2, Fig. 2C**). Similarly, no significant differences in cartilage degeneration were observed between strains 10-wks post-PMX (**Fig. 2D**). Even with clear indications of pain sensing mitigation across models the lack of improved, or worsened, cartilage scores within CD14 KO mice suggests symptomatic OA pain may not be correlated with cartilage degeneration at the time points evaluated in these models.

### Global CD14-deficiency changes the inflammatory cellular landscape within PTOA synovium

We next evaluated synovial histopathology via scoring of common inflammatory features (e.g. lining hyperplasia, sublining cellularity, fibrosis) using a murine specific scoring system on HE stained coronal sections taken midline through the joint (*19*). Histological evaluation revealed no observable strain-related differences in sublining cellularity, lining hyperplasia, or fibrosis scores at 4- or 8-wks post-DMM (**Fig. S2, Fig. 3A**) or 10-wks post-PMX injury (**Fig. 3B**). Overall, little indications of synovitis were observed via histopathology across models at the time points evaluated.

**Fig. 3:**
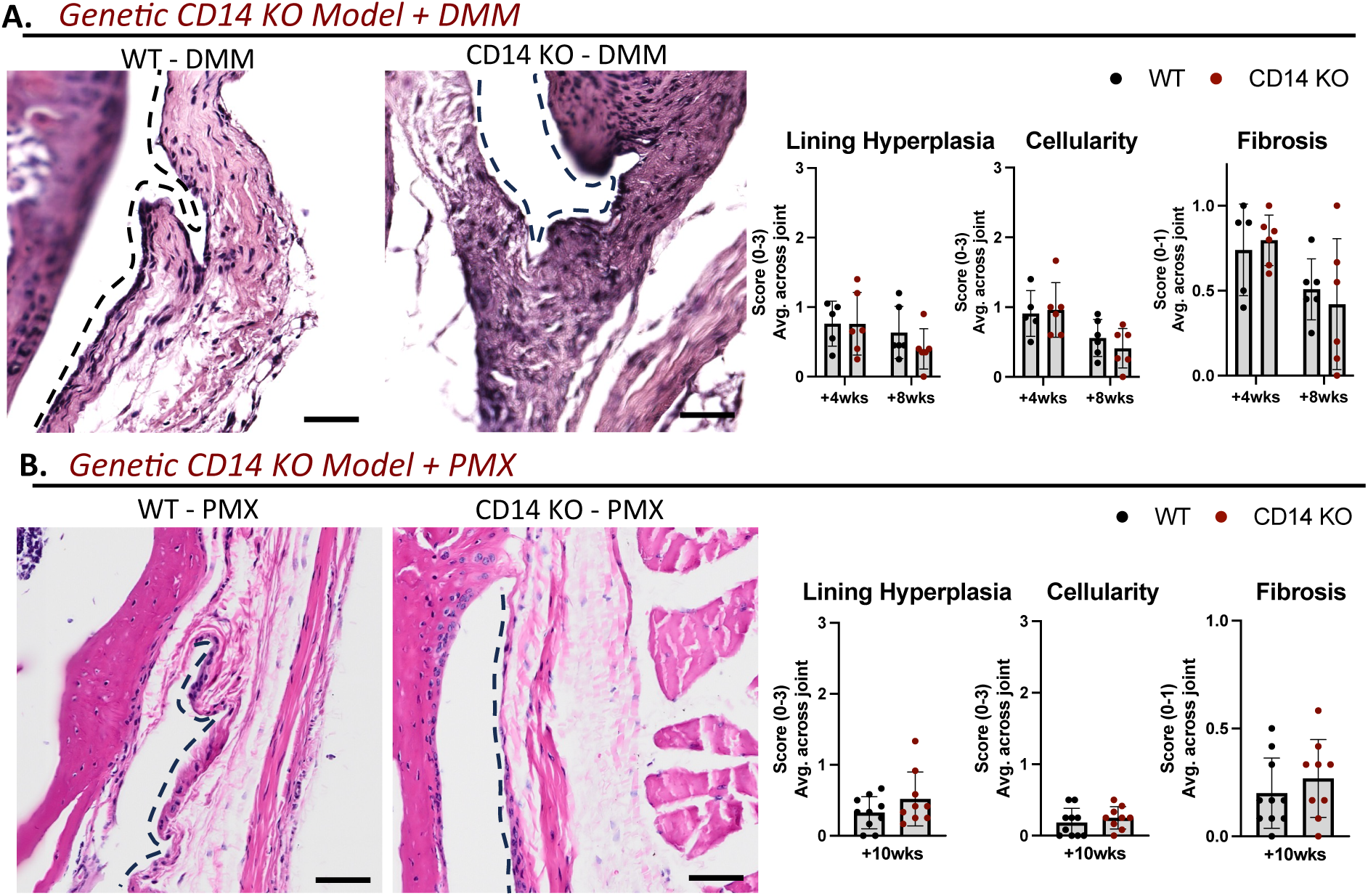
Histopathologic evaluation of synovial pathology following injury. (**A, B**) H&E-stained histological sections from WT (n=6) and CD14 KO (n=6) medial femoral synovial gutters 8-wks following DMM (Scale bar = 50μm) or from WT (n=10) and CD14 KO (n=9) mice10-wks following PMX (Scale bar = 100μm) (Black dashed line indicates the synovium lining). Statistically significant differences were calculated using a student’s T-test with Holm-Šídák post-hoc for multiple comparisons.

We expanded our evaluations via a more in-depth cellular analysis of the synovial inflammatory landscape. First, a basic flow cytometry panel was used to identify synovial leukocyte populations (*CD45+)* including monocytes (*CD45+Ly6C+),* macrophages (*CD45+CD64+),* and T cells (*CD3+, CD3+CD4+)* within pooled synovial tissue (including the fat pad) up to 16 weeks following DMM injury (**Fig. 4A**). Broadly assessing synovial leukocyte populations, no significant changes in the % of *CD45+* cells (of the total synovial cell population) were observed across strains or post-DMM **(Fig. 4B**). However, DMM significantly increased the number of *CD45+* cells within WT synovium at 4-wks post-surgery (p=0.0009), compared to baseline (non-operated) controls. Strikingly, no significant differences in the proportion or number of synovial *CD45+* cells was observed between the CD14 KO and WT strains post-DMM compared to baseline controls (**Fig. 4B**). DMM produced an increase in the % of monocytes (*Ly6C+CD64*-, expressed as a proportion of the *CD45+* population) in WT mice at both 4-wks (p=0.011) and 8-wks (p=0.0002) compared to baseline. In CD14 KO mice, this metric was significant only at 8-wks post-DMM (p=0.0057) (**Fig. 4B**). In WT mice DMM increased the total number of monocytes at 4-wks (p=0.0001) (**Fig. 4B**), while no change in the total number of monocytes was observed post-DMM in CD14 KO mice (**Fig. 4B**). The % and number of macrophages (*Ly6C-CD64+)* significantly increased from baseline in both WT (p<0.0001) and CD14 KO (p<0.005) synovium at 4-wks, but the proportional increases were sustained at 8-wks only in WT animals (p=0.0003) (**Fig. 4B**). Again, the % and total number of cells positive for both *Ly6C* and *CD64* were significantly increased in both WT (p<0.0001) and CD14 KO (p<0.0001) synovium at 4-wks compared to their respective baseline values (**Fig. 4B**). However, the total number of *CD45+Ly6C+CD64+* cells was significantly increased in WT compared to CD14 KO synovium (p=0.0004) at 4-wks post-DMM (**Fig. 4B**). Flow cytometry revealed DMM to produce a clear peak in the total number of monocytes and macrophages 4-wks post-surgery within WT synovial tissue. CD14 KO effectively mitigated this increase, providing initial evidence of CD14 broadly influencing the synovial inflammatory response during PTOA progression.

**Fig. 4:**
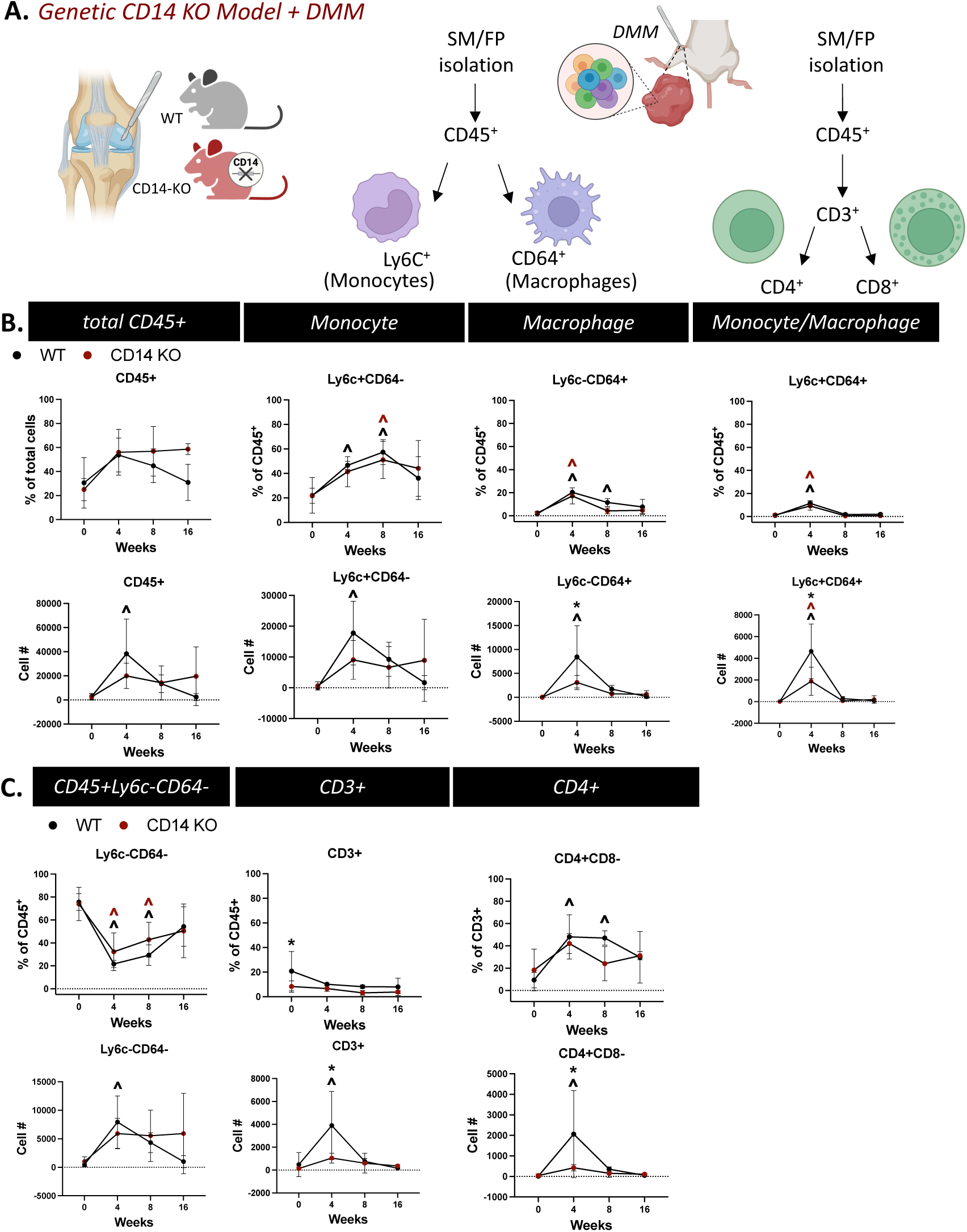
Flow cytometry analysis of synovial immune cell landscape following DMM injury. (**A**) Schematic of flow cytometry analysis for monocytes, macrophages, and T cell populations within DMM synovium and fat pad tissue from WT (n=4-8) and CD14 KO (n=3-5) mice. (**B**) Presence of leukocyte (*CD45+*), monocyte (*Ly6C+CD64-*), and macrophage (*Ly6C-CD64+*, *Ly6C+CD64+*) cell populations represented as cell proportions (%) or as the total number of cells in synovial/fat pad tissue digests. (**C**) Presence of non-monocytic leukocyte (*CD45+Ly6C-CD64-*), T Cell (*CD3+*, *CD3+CD4-CD8-*), and helper T Cell (*CD3+CD4+CD8-*) populations represented as cell proportions (%) or as the total number of cells within synovial/fat pad tissue digests. * Indicates p<0.05 significance between WT and CD14 KO groups at a given time point, ^ indicates p<0.05 significance between 0-wks and 4-, 8-, or 16-wks time points within groups, with color indicating group (i.e. ^: 0-wk WT vs. 4-wk WT). Statistically significant differences were calculated using a use two-factor (factorial) ANOVA followed by Šídák post-hoc for multiple comparisons.

We also observed dynamic shifts in *CD45+* cell populations that were negative for the monocyte/ macrophage markers. Specifically, both CD14 KO and WT synovial tissue had a significantly decreased % of *CD45+Ly6C-CD64-*cells (expressed as proportion of the total *CD45+* population) at 4- and 8-wks post-DMM compared to their respective baseline (non-operated) controls (**Fig. 4C**), while only WT synovium had significantly increased overall number of *Ly6C-CD64-*cells at 4-wks post-DMM compared to WT pre-operative levels (p=0.0012) (**Fig. 4C**).

To further probe this leukocyte cell population, we expanded our evaluations to adaptive immunity and evaluated T cell populations. The % of *CD3+* T cells was significantly higher in WT compared to CD14 KO synovium at baseline (p=0.019, **Fig. 4C**). At 4-wks, DMM produced an increase in the total number of *CD3+* T cells in WT mice compared to baseline (p<0.0001), but not in CD14 KO mice (**Fig. 4C**). Further, at this 4-wk time point CD14 KO mice had significantly lower *CD3+* T cells compared to WT mice (p=0.0021) (**Fig. 4C**). DMM significantly increased helper T cell %’s (*CD3+CD4+CD8-*) in WT mice at 4-(p=0.0015) and 8-wks (p=0.0008) compared to baseline (**Fig. 4C**). Likewise, DMM produced an increase in the total number of helper T cells in WT synovium at 4-wks compared to baseline (p<0.0001) (**Fig. 4C**). While CD14 KO synovium had significantly lower total number of helper T cells compared to WT at 4-wks post-DMM (p=0.0043, **Fig. 4C**). Similar to the evaluations of monocyte/macrophages, CD14 KO mitigated the DMM induced increase in T cells, suggesting CD14 to broadly influence the recruitment of leukocytes to the synovium during OA.

We next employed single-cell RNA-sequencing (scRNA-seq) to assess the synovial immune cell landscape at a higher resolution, focusing on the 4-wk post-DMM time point that showed the most pronounced differences between strains by flow cytometry. Synovial tissue was pooled from 15 knees each of male WT and CD14 KO mice 4-wks following DMM surgery. Clustering of this scRNA-seq data revealed 13 cell clusters with distinct transcriptional profiles, which consisted primarily of varied monocyte, macrophage, and fibroblast populations (**Fig. 5A, B**). Consistent with results from flow cytometry, CD14 KO synovium had substantially decreased populations of a classical monocyte (*Cd45+, Fcgr1/Cd64-, Acod1+, S100a8+, S100a9+, Smox+, Ccl3*+) (*20–22*) and a non-classical interferon activated monocyte population (*Cd45+, Fcgr1/Cd64-, Fcgr3/Cd16+, Ifit1+, Rsad2+, Isg15+)* (*23*) compared to WT synovium (**Fig. 5A,B**). To better identify monocyte subpopulations influenced by CD14-deficiency, classical and nonclassical monocyte populations were re-clustered. Four distinct subclusters were identified and characterized by differential expression of interferon induced transmembrane protein 1 (*Iftm1*), C-C motif chemokine ligand 3 (*Ccl3*), resistin like gamma (*RetlngI*), and interferon regulatory factor (*Irf7*). To gain insight into the function or origin of these clusters, we performed pathway analysis of differentially expressed gene sets within each subcluster (Reactome.org) (*24*). The *Iftm1*+ monocyte population showed over-representation of pathways indicative of “nerve growth factor (NGF) -stimulated transcription” (Reactome pathway identifier: R-HSA-9031628) and “signaling by neurotrophic receptor tyrosine kinase 1 (NTRK1)” (R-HSA-187037), (**Fig. 5C-E**, **Data file S1**). The *Ccl3*+ monocyte population demonstrated overrepresentation of pathways associated with general inflammatory activation, including interleukin stimulation and cell recruitment pathways (R-HSA-449147 & R-HSA-9664424). Interestingly, the *Retlng*+ monocyte population showed over-representation of pathways pointing to TLR stimulation or deficiency (MyD88 deficiency: R-HSA-5602498, IRAK4 deficiency: R-HSA-5603041) (**Fig. 5C-E**, **Data file S1**). Lastly the Irf7+ population had clear over-representation of interferon (IFN)-signaling related pathways (R-HSA-877300, R-HSA-909733) (**Fig. 5C-E**, **Data file S1**). Notably, all 4 re-clustered monocyte subpopulations were decreased in terms of absolute cell number in CD14 KO synovium compared to WT synovium at this timepoint (**Fig. 5C-E**). These findings further support the flow cytometry analyses indicating a broad inflammatory mitigation within leukocyte populations, specifically monocytes, within CD14 KO PTOA synovium.

**Fig. 5:**
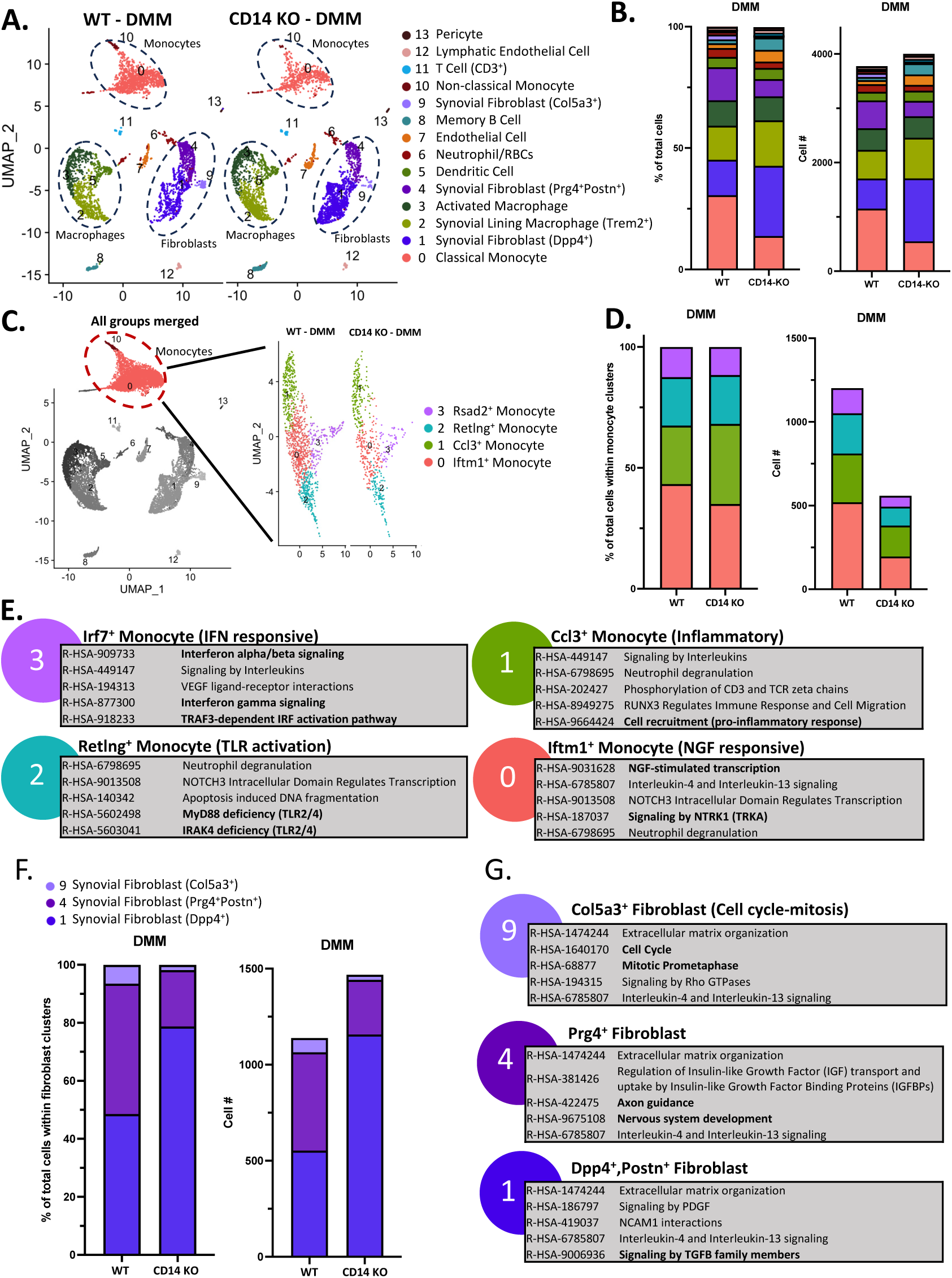
scRNA-seq of synovial tissue 4-wks following DMM injury. (**A**) UMAP and cell cluster assignment across WT and CD14 KO synovium harvested 4-wks post-injury (pooled n=15/group). (**B**) Cell cluster breakdown represented as the percentage of total and number of cells. (**C**) UMAP of re-clustered monocyte populations. (**D**) Breakdown of re-clustered monocytes represented as the percentage of total and number of cells. (**E)** Pathway analysis of re-clustered monocyte groups (Reactome.com) using the top differentially expressed genes. (**F**) Fibroblast cluster breakdown represented as the percentage of total and number of cells. (**G)** Pathway analysis of fibroblast clusters (Reactome.com) using the top differentially expressed genes.

Consistent with prior studies of immune cells within OA synovium (*2*), we identified substantial populations of activated macrophages (*Fcgr1*+), synovial lining macrophages (*Trem2*+, *Adgre1*+), and dendritic cells (*H2-Ab1*+, *Cd209a*+, *Cd74*+) (**Fig. 5A, B**). We also observed smaller populations of neutrophils (*Ly6g*+, *Cd177*+, *Ngp*+, *Camp*+), memory B cells (*Igkc*+, *Cd79a*+, *Bank1*+), and T cells (*Cd3*+, *Thy1*+) (**Fig. 5A, B**). In contrast to monocytes, these additional immune cell populations were mostly similar between strains post-DMM (**Fig. 5A, B**). Lastly, we identified unique *collagen type V alpha 3 chain+* (*Col5a3+*), *proteoglycan 4*+ (*Prg4*+), and *dipeptidyl peptidase 4+* (*Dpp4*+) fibroblast populations across WT and CD14 KO synovium. Interestingly, the *Dpp4*+ fibroblast population increased in the CD14 KO synovium compared to WT, while both the *Col5a3+* and *Prg4*+ fibroblasts decreased in the CD14 KO synovium compared to WT (**Fig. 5F,G**). Pathway analysis suggested that the *Dpp4*+ fibroblasts, which had a greater presence in CD14 KO synovium, has over-representation of TGFB signaling pathways (R-HSA-9006936) (Data file S2). *Prg4*+ fibroblasts, which were more predominant in the WT synovium, exhibited over-representation of pathways involved in axon guidance (R-HSA-422475) and nervous system development (R-HSA-9675108), (**Fig. 5F,G**, **Data file S2**). Lastly, the smaller *Col5a3+* sub-population which had a greater presence in the WT synovium, was likely a replicating fibroblast, with over-representation of cell cycle (R-HSA-1640170) and mitosis related pathways (R-HSA-68877) (**Fig. 5F,G**, **Data file S2**). Sequencing results indicate CD14 KO to clearly mitigate monocyte activity while also possibly biasing the synovial landscape away from neuro-promoting fibroblasts, which may relate to the mitigated pain responses within CD14 KO mice.

We next employed spatial proteomics to extend our analysis beyond the transcriptome and further probe the synovial cellular landscape 4-wks following DMM injury. Utilizing imaging mass cytometry (IMC) we applied a panel of 24 metal tagged antibodies (**Table S2**) to sagittal knee sections taken from the medial side of the knee. IMC identified 14 unique cell clusters with varying protein expression across our panel (**Fig. 6A**) (*25*). This included clusters with protein signatures of non-immune cell populations such as adipocytes (Cluster 1, perilipin), vascular cell types (Cluster 8, CD31^high^), and fibroblasts (Clusters 5 & 6, αSMA+), as well as clusters of immune cell populations with signatures of neutrophils (Cluster 7, Ly6G^high^), and macrophages (Clusters 9&10, F4/80^high^). Two pan-positive hematopoietic cell clusters (CD45+) were identified and separated by the presence of the proliferative marker Ki67 (Clusters 10 & 14). Lastly, multiple CX3CR1+ populations were observed, categorized as monocytes (Cluster 12, F4/80^low^) and resident lining macrophages (Clusters 13: F4/80^high^) (**Fig. 6A**) (*26*).

**Fig. 6:**
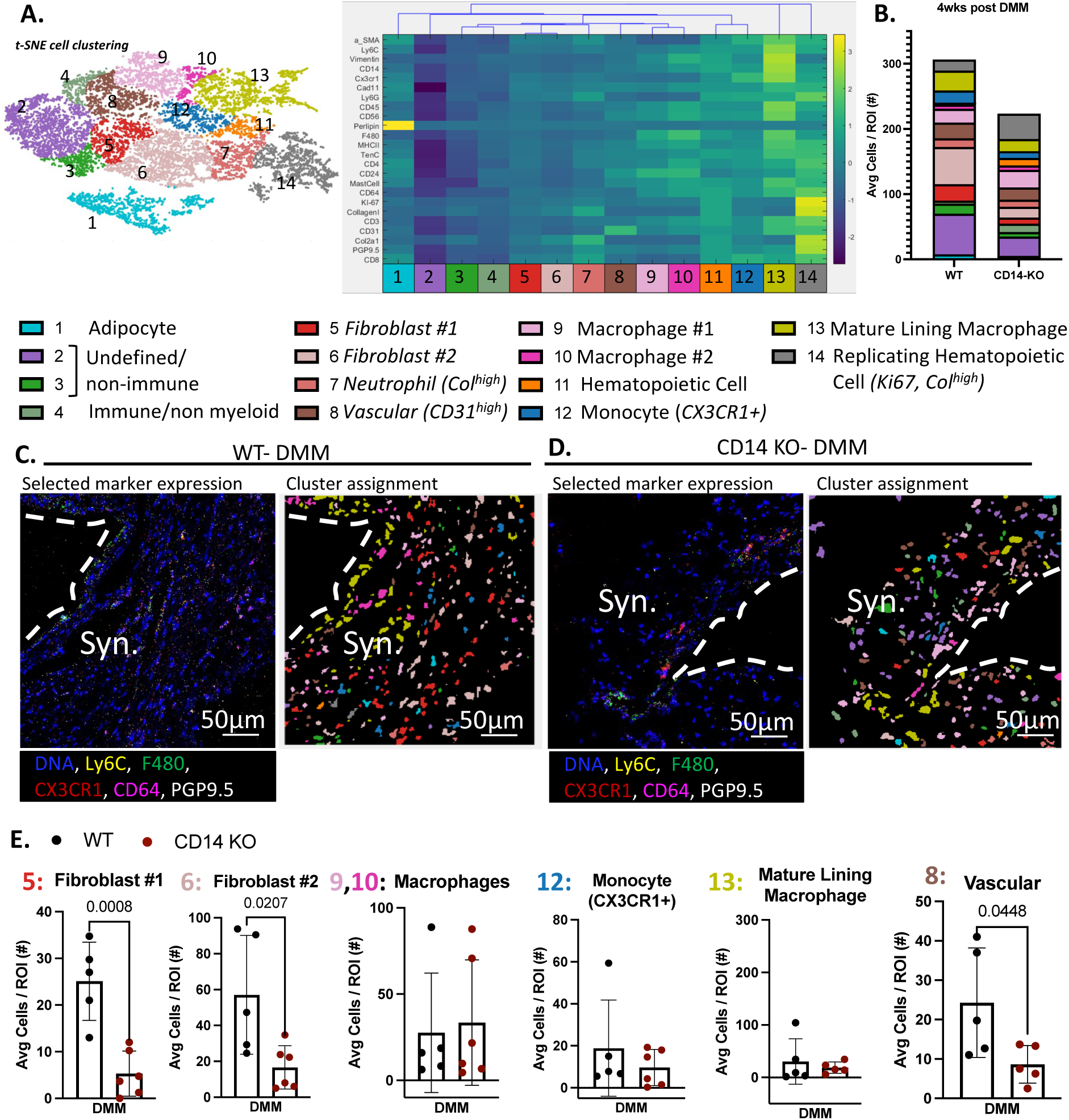
IMC analysis of synovial cell phenotypes 4-wks following DMM injury. (**A**) Cell cluster t-SNE scatter plot with normalized heatmap of protein panel marker expression and cluster assignments. (**B**) Cell cluster breakdown represented as the average number of cells within each synovial ROIs per sample (n=5-6 samples/genotype). (**C, D**) Generated map of selected marker expression and associated cell cluster assignment within WT and CD14 KO synovial ROIs 4-wks post-DMM. Synovial lining = white-dashed line. (**E**) Cell number analysis represented as the average number of cells within synovial ROIs. Statistically significant differences were calculated using a student’s T-test.

IMC analysis revealed strain-related differences in the cellular landscape of the synovium, particularly among synovial fibroblasts (**Fig. 6A-D**). Quantification revealed significant reductions in two synovial fibroblast clusters (Cluster 5, p<0.0001 & 6, p=0.0020) in CD14 KO compared to WT synovium (**Fig. 6E**), consistent with the scRNA-seq dataset. Quantification of macrophage populations (Clusters 9&10), including the lining macrophage population (Cluster 13), revealed no significant changes in cell numbers between WT and CD14 KO synovium (**Fig. 6E**), consistent with our scRNA-seq (**Fig. 5**). While a possible monocyte population could be observed, no clear significant changes in cell numbers were seen between WT and CD14 KO synovium (**Fig. 6E**) using this approach. Further, WT synovium showed an increase in the CD31^high^ vascular cell cluster (p=0.045, Cluster 8), compared to the CD14 KO synovium (**Fig. 6E**). Harnessing the spatial information gained via IMC, we performed a neighborhood analysis, revealing CD14 KO to have no effect on the number of neighboring cell cluster interactions within the DMM synovium, with no significant changes compared to WT (**Fig. S3**). Though analyses are restricted to the synovial gutter regions, spatial proteomic data provided by IMC further revealed how CD14 KO influences the PTOA synovial landscape. Suggesting CD14 KO to modulate PTOA synovial fibroblast populations, also observed within scRNA-seq analysis, in addition to possibly mitigating vascularity within the synovium.

### Local delivery of a CD14 neutralizing antibody mitigates spontaneous and evoked pain behaviors across mild and severe PTOA models

To add to the translatability of our results, we evaluated the efficacy of local pharmacologic CD14 blockade on OA pain response and disease progression. In addition to the surgical DMM model, to eliminate the effects of the surgical intervention we compared responses to a non-surgical load-induced severe model of anterior cruciate ligament rupture (ACLR) (*25*). To further challenge this approach, the ACLR procedure was performed on older (36-wk old) female mice that were fed a high fat diet (ACLR-HFD) from 16-wks of age to induce obesity and incorporate metabolic stress into the model. In both models, mice were treated with intra-articular injection of either a monoclonal anti-murine CD14 antibody or a murine IgG2a isotype control (both 0.5mg/kg). Three weekly injections beginning 48 hours post-DMM or ACLR-HFD injury were performed.

In the DMM model, we saw the expected post-DMM decreases in knee withdrawal threshold in IgG treated mice 4-wks post-DMM injury (p=0.0078), indicating development of knee hyperalgesia (**Fig. 7A**). No significant post-DMM change in withdrawal threshold was observed at 4-wks in the anti-CD14 treated group (**Fig. 7A**). Spontaneous cage behavior and overall mobility was evaluated using the Laboratory Animal Behavior Observation Registration and Analysis System (LABORAS^TM^, Metris) (*11*); activities (e.g. climbing, rearing, grooming) and mobility metrics (e.g. total distance traveled) were measured over a 14-hr testing period. Mice that received the anti-CD14 antibody spent more time rearing at 4-(p=0.0052) and 8-wks (p=0.011) post-DMM when compared to IgG treated mice and compared to their pre-surgery levels (p=0.0006 at 4 wks; p<0.0001 at 8 wks) (**Fig. 7A**). Average speed (4- & 8-wks), and total distance traveled (4- & 8-wks) were increased in anti-CD14 treated mice compared to IgG treated mice post-DMM (**Fig. S4**). No changes to eating or drinking behavior or weight were observed between IgG and anti-CD14 treated mice at pre-operative or post-DMM time points (**Fig. S4**). DMM injury increased front to rear paw weight bearing ratios in IgG treated mice at 4-(p=0.032) and 8-wks (p=0.012), while no significant changes in front to rear paw weight bearing ratios were observed in anti-CD14 treated mice (**Fig. 7A**). No changes in left to right rear paw weight bearing ratios up to 8-wks post-DMM were observed in either treatment group (**Fig. 7A**). We next tested whether delaying treatment after DMM injury could still be effective. Again, male WT mice underwent surgery and 30 days later were given their first intra-articular injection of the IgG control or anti-CD14 antibody, followed by weekly injections for the next two weeks (**Fig. S5**). Interestingly, no difference in spontaneous cage behavior or paw weight distribution was observed between groups when treatment was delayed (**Fig. S5**). The mitigation of evoked and spontaneous pain behaviors within the DMM model largely mimicked the results of the global genetic knockout, while the time of delivery (early vs. late) dependence suggests CD14 to play a more substantial role in the early inflammatory and disease initiating events within the first 4-wks following injury.

**Fig. 7:**
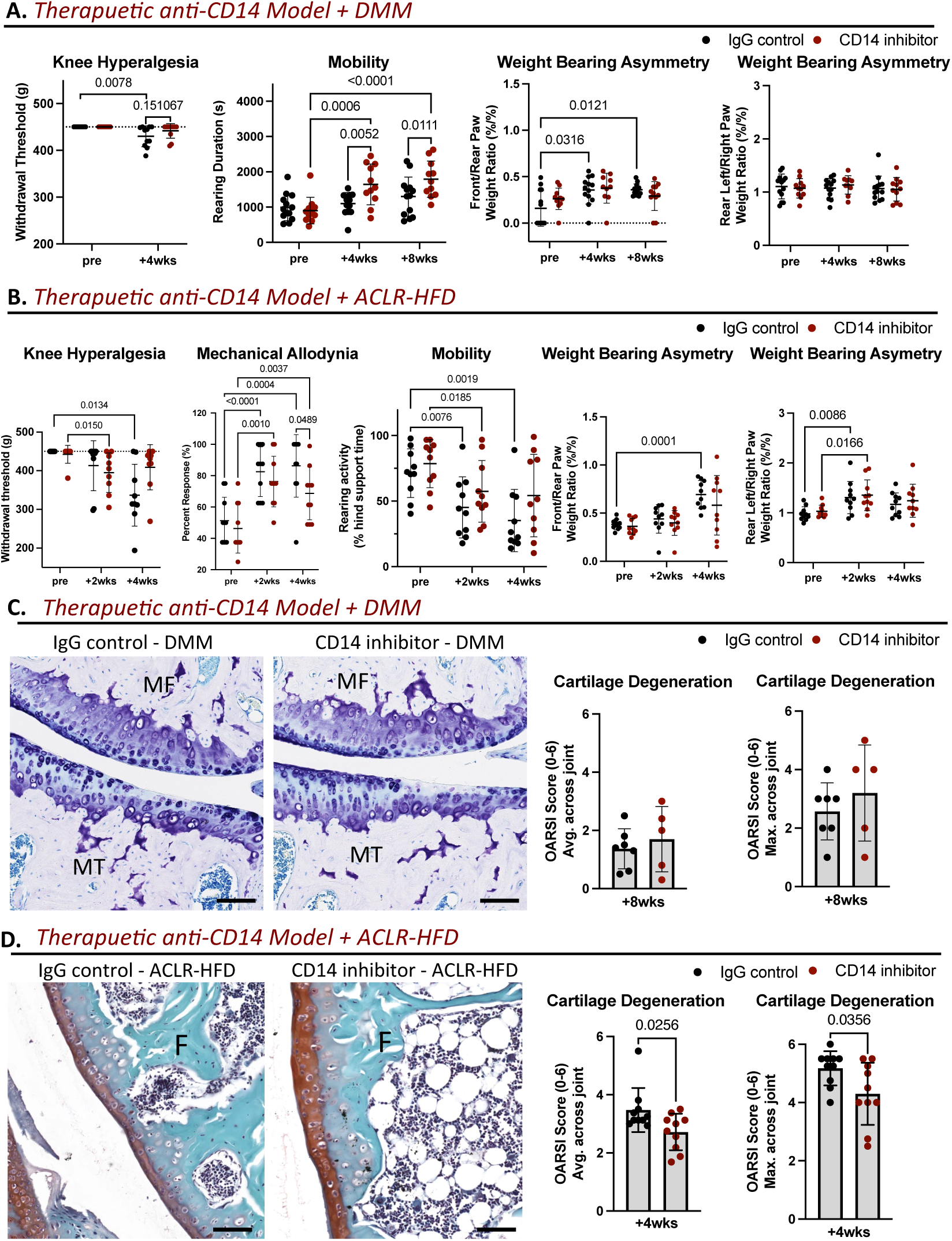
Effect of blocking CD14 via neutralizing antibody on pain and cartilage damage following injury. (**A**) Pain, mobility, and weight bearing changes between IgG (n=11) or anti-CD14 (n=11-12) treated WT. Knee hyperalgesia measured via withdrawal threshold (g) using a PAM device. Mobility measured via rearing duration (s) using the LABORAS system. Weight bearing asymmetry measured as the front to rear paw weight ratio (%/%) and rear-left to rear-right paw weight ratio (%/%). (**B**) Pain and weight bearing changes between IgG (n=9-10) or anti-CD14 (n=9-10) treated WT mice. Knee hyperalgesia measured via withdrawal threshold (g) using a PAM device. Mechanical allodynia measured via the 50% withdrawal threshold (g) using von Frey fibers. Weight bearing asymmetry measured as the front to rear paw weight ratio (%/%) and rear-left to rear-right paw weight ratio (%/%). (**C**) Toluidine Blue stained histological sections of IgG (n=7) and anti-CD14 (n=5) treated knees 8-wks following DMM (Scale bar = 100μm). (**D**) Safranin-O-stained histological sections of IgG (n=10) and CD14 inhibitor (n=10) treated knees 4-wks following ACLR-HFD (Scale bar = 100μm). Statistically significant differences between treatment groups in knee hyperalgesia (**A,B**) were calculated using the Mann-Whitney U non-parametric T-test with Holm-Šídák post-hoc, and across time points using either the Friedman test with Dunn’s multiple comparison adjustment or the Kruskal-Wallis test with Dunn’s multiple comparison adjustment for data sets with missing values. Statistically significant differences in remaining behavioral outcomes (**A,B**: mobility, allodynia, & weight bearing asymmetry) were calculated using a use repeated measures mixed effect model followed by Šídák post-hoc for multiple comparisons. Statistically significant differences in histopathological scoring (**C,D**) were calculated using a student’s T-test.

In the more severe ACLR-HFD model, control IgG treated mice developed significant hyperalgesia at 4-wks post-injury (p=0.013) compared to preoperative measurements (**Fig. 7B**). Anti-CD14 treated mice saw more transient effects, with a significant increase in hyperalgesia by 2-wks post-injury (p=0.015 vs. preoperative levels), which resolved by the 4-wk post-injury time point (**Fig. 7B**). Mechanical allodynia developed following ACLR-HFD in both groups at both 2- and 4-wks (**Fig. 7B**). However, anti-CD14 treatment significantly reduced allodynia (% response) at 4-wks post-ACLR-HFD compared to IgG treatment (**Fig. 7B**). ACLR-HFD significantly decreased rearing activity in both IgG (p=0.0076) and CD14 blockade (p=0.019) treated mice at 2-wks, when compared to preoperative values (**Fig. 7B**). This decrease in rearing activity was sustained up to 4-wks post-ACLR-HFD in IgG (p=0.0019) but not CD14 blockade treated mice (**Fig. 7B**). ACLR-HFD produced a significant increase in the front to rear paw weight bearing ratio in the IgG treated group at 4-wks (p=0.0001), with no significant change observed in anti-CD14 treated mice (**Fig. 7B**). Further, ACLR-HFD produced significant increases in rear left to rear right paw weight bearing ratios at 2-wks, in both IgG (p=0.0086) and anti-CD14 (p=0.0086) treated mice (**Fig. 7B**), with no significant differences between treatment groups. Although changes in these outcome parameters were of greater magnitude in this more severe model, overall results with early anti-CD14 treatment were similar to that observed in the milder, DMM model of PTOA.

### Local pharmacologic blockade of CD14 provides mild protection against histopathologic cartilage degeneration and synovial fibrosis in the ACLR-HFD model

We next evaluated cartilage damage and synovial pathology in both the DMM and ACLR-HFD models. Cartilage degeneration was neither improved nor worsened with delivery of anti-CD14 in the DMM model, with similar average and max OARSI scores at 8-wks across IgG and anti-CD14 treated groups. This was consistent in both the early (**Fig. 7C**) and delayed treatment regimens (**Fig. S5**). In both groups mild losses to proteoglycan staining, surface fibrillation, and loss of non-calcified cartilage tissue were seen (**Fig. 7C**). In contrast, in the more severe ACLR-HFD model, significant but modest reductions in both average (p=0.036) and max (p=0.026) OARSI scores were observed in anti-CD14 treatment mice compared to IgG treatment at 4-wks post-injury (**Fig. 7D**). The most notable effect was increase in the preservation of proteoglycan staining in the femoral cartilage surface in CD14 blockade treated mice compared to IgG-treated mice (**Fig. 7D**).

Synovial pathology was then evaluated in both models. In the DMM model, a mild but non-significant reduction (p=0.07) in sublining cellularity was observed in the anti-CD14 treated compared to IgG treated knees at 8-wks post-surgery (**Fig. 8A**). No significant changes were observed in lining hyperplasia or fibrosis between treatment groups (**Fig. 8A**). In the ACLR-HFD model, anti-CD14 treatment produced a mild but non-significant decrease (p=0.074) in fibrosis scores at 4-wks post-injury when compared to IgG treated mice, with no differences in lining hyperplasia or cellularity observed between treatment groups (**Fig. 8B**). Ultimately, CD14 blockade produced minimal changes to tissue histopathology at early-stage disease progression, similar to what was observed within the global CD14 KO models.

**Fig. 8:**
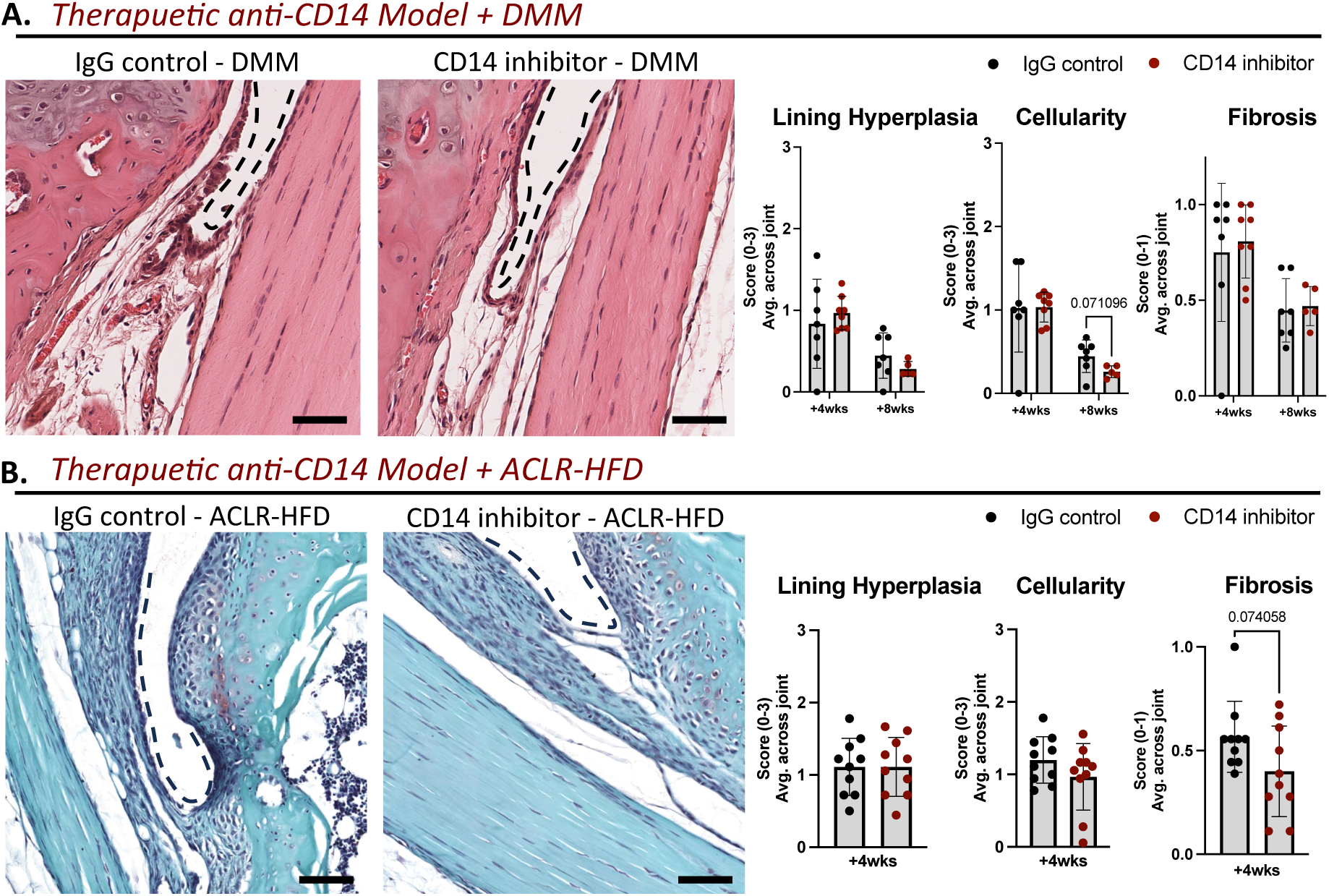
Effect of CD14 blockade on synovitis following injury. (**A**) H&E-stained histological sections of IgG (n=7) or CD14 inhibitor (n=5-8) treated knee joints 8-wks following DMM (Scale bar = 50μm). (**B**) Safranin-O-stained histological sections of IgG (n=10) or CD14 inhibitor (n=10) treated knee joints 4-wks following ACLR-HFD (Scale bar = 50μm) (Black dashed line indicates the synovium lining). Statistically significant differences were calculated using a student’s T-test with Holm-Šídák post-hoc for multiple comparisons.

## DISCUSSION

CD14 is of interest as a potential biomarker of OA severity and synovial inflammation, as synovial fluid levels of the soluble form are associated with disease progression, symptoms, and synovial macrophage infiltration (*4–7*). These clinical observations, and the known inflammatory function of CD14, indicate a possible pivotal role of CD14 in facilitating synovial inflammation and influencing disease. While OA pain is frequently localized to the joint and exacerbated during movement, many patients exhibit a more persistent, chronic, pain phenotype, and recent studies have begun to identify a multitude of OA pain phenotypes. This complexity of pain within the human OA population is one factor contributing to the low correlation between pain severity and structural damage in this disease (*27, 28*). Moreover, common comorbidities and risk factors such as obesity can further complicate the relationship between pain, function, and joint pathology (*29*). Given this complexity within the human OA population, novel approaches for mitigating OA symptoms need to be tested across a range of OA disease models and incorporate common comorbidities in order to identify the most promising methods for future success in translation.

With this in mind, we employed models of PTOA that captured a range of mild (DMM) to more severe (PMX, ACLR-HFD) pathology and evoked pain responses. For example, robust knee hyperalgesia was observed within the PMX model of PTOA, while in mid-stage DMM milder knee hyperalgesia was present. Across these models, we tested effects of genetic deficiency of CD14 as well as antibody blockade via intra-articular delivery. Our results demonstrated that both approaches consistently improved both evoked and spontaneous pain behaviors across models (**Table 1**). Specifically, global CD14-deficiency was protective against increased evoked pain responses in both the mild DMM and more severe PMX models, in addition to improving spontaneous locomotive behavior post-DMM and ameliorating weight bearing asymmetry post-PMX. Adding to the variability of human pain phenotypes, females are known to have a higher prevalence of OA, with more severe pain (*30*). Incorporating female mice in the more severe PMX and ACLR-HFD models, we saw similar CD14-mediated protections against painful behavior. Ultimately these results provide strong support that the targeting of CD14 could offer broad protection from pain and functional deficits after OA-inducing joint injuries in both males and females.

**Table 1:** Summary of outcomes, and time point at which an effect of CD14 modulation was seen across models. Dash. (**-**) **denotes the outcome was not performed in a specific model.**

|  | Global CD14 KO | Global CD14 KO | CD14 blockade | CD14 blockade |
| --- | --- | --- | --- | --- |
| *Outcome* | DMM | PMX | DMM | ACLR-HFD |
| *Knee Hyperalgesia* | Mitigation (4- & 8-wks) | Mitigation (4- & 8-wks) | Mild Mitigation (4- & 8-wks) | Mild Mitigation (4-wks) |
| *Mechanical Allodynia* | - | Mitigation (4-wks) | - | Mitigation (4-wks) |
| *Cage Behavior/Mobility* | Improved (0-4-wks) | - | Improved (0-4,8-wks) | Mild Improvement (4-wks) |
| *Weight Bearing Asymmetry* | No protection | Mitigation (10-wks) | Mitigation (4,8-wks) | Mitigation (4-wks) |
| *Cartilage Degeneration Score* | No protection | No protection | No protection | Protection (4-wks) |
| *Synovitis Score* | No protection | No protection | No protection | No protection |

It is thought that cartilage damage leads to symptomatic joint pain, yet the two are not well correlated (*31*). Our evaluation of therapeutic CD14 mitigation on cartilage degeneration in these models supports this idea, with disparate effects on pain and cartilage pathology. Global deficiency of CD14 provided consistent protection in pain and functional outcomes, yet no clear protection from post-injury cartilage degeneration within the timeframes evaluated (8-wks post-DMM, 10-wks post-PMX). We previously reported protection from cartilage degeneration in the CD14 KO strain, but this was observed only later, at 19-wks post-DMM, a time point not examined in this study (*11*). In the current study, we did observe protection from cartilage degeneration following anti-CD14 treatment within the more severe ACLR-HFD model, which results in rapid development of a more severe pathology than seen in the DMM model. These results demonstrate that the impact of CD14 on PTOA pain is temporally distinct from its impact on cartilage pathology, with relevance to pain mechanisms at earlier stages of disease, while the impact on cartilage pathology might not be seen until more severe pathology has developed.

Of note, neither genetic deficiency nor pharmacologic blockade worsened joint pathology. This is important as other anti-inflammatory approaches in PTOA models have been shown to potentially worsen joint tissue damage, presumably by impacting the protective effects of resident immune cells, such as macrophages (*32, 33*). Further, the demonstration of the safety and pain mitigating impact of this approach in a non-surgical model (ACLR-HFD), which has high clinical relevance to the initiation of PTOA in humans even in the setting of obesity, is promising. Obesity is both a risk factor for development of OA, and a common comorbidity that can worsen progression of pain and structural damage (*29*).

As synovial pathology has been linked to joint pain in OA, we hypothesized that effects of CD14-deficiency and blockade might be mediated through modulation of synovial inflammation. Our initial histopathologic evaluation of synovial inflammation revealed mild synovial pathology in all three models, consistent with prior studies (*19, 25*), with little modulation by genetic or therapeutic targeting of CD14 when evaluated at this length scale. We next employed more robust approaches to phenotype the cellular landscape within the synovium. Flow cytometry initially revealed CD14-deficiency to clearly mitigate monocyte, macrophage, and T cell populations within the DMM synovium. scRNA-seq further revealed a broad decrease in all identified monocyte subpopulations within CD14 deficient DMM synovium. Evaluating these monocyte subpopulations, pathway analysis suggested inflammatory activation by different mechanisms, at least two which are downstream of TLRs. Specifically, the *Retlng+* “TLR responsive” monocyte population exhibited over representation of MyD88 and IRAK4 pathway activation and upregulated expression of the common TLR4 activating DAMPs *S100a8* and *S100a9* (*34*). The *Irf7+* “IFN responsive” monocyte population exhibited over representation of interferon alpha/beta signaling and IRF pathways, and upregulation of a host of broad IFN response genes. Notably *Irf7* is a transcription factor essential for IFN-α/β gene expression, and is activated downstream of TLRs primarily by the MyD88 independent pathway (*35*). The CD14 dependent decreases in these two inflammatory monocyte subpopulations is consistent with its known role as a pattern-recognition receptor, recognizing various DAMPs and PAMPs and facilitating TLR activation. Moreover, CD14 is responsible for internalization of TLR4, which is required to trigger the MyD88-independent activation of *Irfs* and production of type I interferons (*36*). These high dimensional evaluations of the synovial inflammatory landscape are consistent with the known role of CD14 in myeloid lineage cell inflammatory responses, and highlight the limitations of relying solely on synovial histopathology to detect meaningful inflammatory changes in the synovium within these models (*6, 7*).

scRNA-seq and pathway analysis also revealed a CD14-mediated decrease in a *Iftm1+* monocyte population, which we categorized as possibly “NGF-responsive” based on its increased expression of *Egr3*. *Egr3* is a transcriptional regulator downstream of NGF stimulation in neurons, while in monocytes NGF stimulation is thought to be involved in enhanced inflammatory activation (*37, 38*). In the context of OA, and coinciding with roles in promoting innervation, NGF has been shown to be an effective target for mitigating OA pain, with clinical trials supporting efficacy of NGF blockade. Unfortunately these results were ultimately complicated by unexpected side effects (*39, 40*) that precluded their ultimate clinical translation. However, following the success of anti-NGF treatment in mitigating OA pain, it is possible that the decreased number of inflammatory activated monocytes within the CD14 deficient synovium may limit NGF driven neuroinflammatory pathways, and contribute to the protections demonstrated in this study. Further studies are needed to pinpoint possible mechanisms by which the CD14-mediated decreases in these synovial monocyte subsets during OA development relates to the mitigation of pain.

Beyond the CD14-mediated decrease in synovial monocyte populations, we also observed a dynamic change in synovial fibroblast subsets following DMM. IMC identified significant decreases in two possible fibroblast clusters within CD14-deficient DMM synovium, defined by their positive expression of αSMA, Col1a, and Col2a1, and low expression of CD45 and myeloid panel markers. Likewise, scRNA-seq indicated decreases in two synovial fibroblast subsets in the CD14-deficient synovium post-DMM, including a *Prg4+* fibroblast population with over-representation of transcriptional pathways involved in “axon guidance” and “nervous system development” and a smaller *Col5a3*+ fibroblast population. Though further work is needed to demonstrate whether either subset is actively involved in changes in innervation within the synovium observed in OA models (*41*), others have demonstrated that synovial lining fibroblast subsets from RA patients are specifically associated with pain and sensory nerve growth (*42*).

Our scRNA-seq results also revealed an increased *Dpp4+, POSTN+* fibroblast subset within CD14 deficient DMM synovium, hallmarked by over-representation of TGF beta signaling pathway related genes. This suggests a potential pathologic role in synovial tissue fibrosis (*43*). It is reassuring that our histopathologic analysis did not show any increase in fibrosis in the CD14-deficient strain. However, more sophisticated methods of evaluating tissue fibrosis may be needed to definitively understand the effect of CD14 on OA synovial matrix composition. Although our IMC analysis did not detect an overall increase in the fibroblast population, it is possible that this population may have been missed by this analysis, which was applied to limited regions of interest in the tissue sections, while the scSeq provided more comprehensive tissue coverage and included synovial and fat pad tissue. Further, IMC utilizes a limited panel of antibodies to define cell populations, while scSeq defines cell subsets based on more comprehensive transcriptional programs, and so results are not exactly replicated by each technique. However, our identification of distinct *Dpp4+* and *Pgr4+* fibroblasts via single cell scRNA-seq in the DMM model is consistent with results published utilizing the ACLR model (*44*). Whether the differences we observed in these populations between strains are a direct result of CD14-deficiency on fibroblasts or an indirect outcome acting through fibroblast crosstalk with nearby CD14-deficient myeloid cells, remains to be tested.

To begin translating these findings, we delivered a monoclonal CD14 neutralizing antibody intra-articularly after injury in the milder DMM model in male mice and the more severe ACLR-HFD model in female mice. In both models, the CD14 neutralizing antibody offered protection against pain and mobility loss, similar to results seen in the setting of genetic CD14-deficiency. Notably, protection was afforded by both the naturally occurring antibody clone (used in the ACLR-HFD model) and the same clone modified to attenuate Fc-receptor binding mediated effects (LALA-PG variant, a common modification made to clinically utilize monoclonal antibodies (*45*) in the DMM model). Importantly, the use of a surgical and a non-surgical model rules out the possibility that results are artifactual and only relevant to surgical injury models. In addition, the finding of significant protection, even in the setting of obesity, and across male and female cohorts, increases the potential of our findings to be translatable. Although only tested in the DMM model, we saw no protective effects with delayed delivery of the anti-CD14 antibody, and so it is possible that therapeutic efficacy may require delivery during the early post-injury period. Of note, a chimeric anti-human CD14 antibody (IC14) delivered systemically has been evaluated in humans in phase 1 clinical trials for the treatment of sepsis (*46*) and neurodegenerative disease (*47*), and in a phase II trial for COVID-19 associated pneumonia (*59*). Although no definitive therapeutic efficacy in these settings was established, these prior studies demonstrate that CD14 antagonism is safe and well-tolerated.

In summary, our results demonstrate that deficiency and pharmacological inhibition of CD14 reduced pain-related behaviors without worsening structural pathology in multiple models of PTOA that range in severity of pain, pathology, progression rates, metabolic stress, and effects on males and females. Moreover, our assessment of the synovial cellular landscape in the DMM model supports that CD14 modulation plays a primary role in attenuating synovial monocyte infiltration and altering fibroblast phenotype shifts following acute traumatic injury. In this cross-institutional study, models were evaluated in three different labs and varied methodologies were employed to detect evoked and spontaneous pain-related outcomes across the three models. The consistent finding of pain-behavior mitigation, regardless of methodology, laboratory location, and model used, adds to the rigor of our analysis and demonstrates reproducibility of our findings. This extensive evaluation supports a primary role of CD14 in mediating pain and mobility loss during PTOA development. Ultimately, our therapeutic findings provide strong support for the future development of this approach in treating painful OA during early disease progression in humans.

## MATERIALS AND METHODS

### Study design

Synovial fluid samples and clinical data were obtained from participants enrolled in on ongoing clinical the MOVE-OK trial (clinicaltrials.gov identifier NCT05035810) the protocol of which has been previously published (**Table S1**) (*12*). A pain pressure threshold (knee hyperalgesia) was measured in clinical trial patients and reported as kg/cm^2^, recorded from the average of three measurements taken at the knee joint using a handheld algometer. Next, the effect of CD14-deficiency on OA progression was evaluated using CD14 KO mice, with C57BL/6 WT mice used as controls (The Jackson Laboratory). Evaluations were carried out in male mice from 4- to 8-wks following DMM surgery, and in male and female mice from 4- to 10-wks post-PMX surgery. Further evaluation of joint inflammation via flow cytometry, scRNA-seq, and spatial proteomics were carried out in CD14 KO and WT mice 4-wks post-DMM surgery. Lastly, the potential of a local CD14 blocking therapeutic was evaluated in both DMM and ACLR-HFD models of PTOA. In the DMM model, an early dose cohort of male WT mice were given 3 weekly doses (0.5mg/kg) of either an anti-CD14 monoclonal antibody or an IgG2a control via intra-articular injection beginning 48-hrs following surgery, while a late dose cohort began treatment 30-days following DMM and continued weekly for a total of 3 doses. DMM mice were evaluated from 4- to 8-wks post-surgery. In the ACLR-HFD model female mice were fed a HFD diet from 16- to 36-wks of age and then underwent non-surgical compressional loading of the knee joint to rupture the ACL at 36-wks of age (*25*). Beginning 48-hrs post injurious loading, mice received 3 weekly doses of either an anti-CD14 monoclonal antibody or an IgG2a control (0.5mg/kg) via intra-articular injection. ACLR-HFD mice were evaluated for 2- to 4-wks post-injury. Additional information is provided in Supplemental materials and methods.

### Pain and behavioral analysis

Longitudinal pain behavioral testing was carried out across DMM, PMX, and ACLR-HDF models. Knee hyperalgesia was measured via the PAM device (Ugo Basile), measuring the withdrawal threshold (g) (*16, 48*). In the PMX model, mechanical allodynia was performed on the ipsilateral (injured) hind paw using von Frey fibers, measuring the withdrawal threshold (median 50% withdrawal) using the up-down method (*16, 17*), while in the ACLR-HFD model the positive withdrawal response % was averaged across a range of von Frey fibers (*49*). Spontaneous behavior was evaluated via measurements of paw weight bearing and cage activity. Paw weight bearing analysis utilized the ADWB (DMM & ACLR-HFD models) and static incapacitance meter systems (PMX model, Bioseb) (*16*), which recorded % of paw weight bearing. The ADWB system was also used to evaluate time spent rearing on the hind legs. In the DMM model only, additional cage behavioral analysis was performed via the LABORAS (Metris), where various cage activities and mobility metrics were measured. Additional information in Supplemental materials and methods.

### Histopathology

Cartilage damage was assessed using the modified Osteoarthritis Research Society International (OARSI) (*18*). Features of synovial inflammation (synovitis) were evaluated using an approach developed for murine models (*19*). Additional information in Supplemental materials and methods.

### Imaging Mass Cytometry (IMC) and cellular phenotype analysis

Paraffin knee joint sections were utilized for IMC analysis (*25*). Sections were stained with a 24-panel metal-conjugated antibody cocktail (**Supplemental Table 1**) and a cationic nucleic acid intercalator to stain for DNA. Stained sections were imaged using the Hyperion Imaging System (Standard BioTools). Raw mass cytometry data was acquired for each 1 μm^2^ pixel within ROIs to generate MCD image files. Single cell masks were created using DNA channel only images (deepcell.org). Cell masks and Multi-OME tiff files for each individual ROI were loaded into IMACytE software, to create t-distributed stochastic neighbor embedding (t-SNE) dimensionality reduction analysis with arcsine transformation to produce data normalization and cluster analysis, as previously described (*25, 50*). Additional information in Supplemental materials and methods.

### Single-cell RNA-sequencing (scRNA-seq)

Synovial single cell suspensions were sequenced by the Children’s Hospital of Philadelphia (CHOP) Center for Applied Genomics (CAG) Sequencing Core. Raw data was pre-processed in Cell Ranger and Uniform Manifold Approximation and Projection (UMAP) mapping was generated with unsupervised clustering using Seurat (R, v4.4.1). Unique gene markers lists were generated to identify clusters. Pathway analysis using Reactome.org was utilized to further identify pathway over representation within clusters (*24*). Additional information in Supplemental materials and methods.

### Statistical analysis

All data shown are shown as means SD. Linear regression analyses was performed in Stata software (v14.2). All remaining statistical analyses, including Friedman’s test for normality, Spearman’s test, Student’s T-test, Mann-Whitney U non-parametric T-test, Kruskal-Wallis, repeated measures mixed effect model, repeated measures two-way ANOVA, Holm-Šídák post-hoc for multiple comparisons, Šídák post-hoc for multiple comparisons, and Dunn’s multiple comparison adjustment, as indicated in figure legends, were performed using Graphpad Prism software (v9). A p-value of <0.05 was considered significant.

## List of Supplementary Materials

Supplemental Materials and Methods

Fig. S1: Pain and weight bearing changes between WT and CD14 KO female mice following PMX injury.

Fig. S2: Histopathologic evaluations of cartilage damage and synovitis at intermediate time points withing CD14 KO and CD14 blockade DMM models.

Fig. S3: Neighborhood analysis of IMC cell clusters across WT and CD14 KO synovium 4-wks post-DMM.

Fig. S4: Behavioral data from early CD14 blockade delivery DMM model.

Fig. S5: Pain, behavioral, and histopathologic evaluations within the late CD14 blockade delivery DMM model.

Table S1: Synovial fluid patient characteristic information.

Table S2: List of antibodies used for IMC.

Data file S1: Monocyte cluster pathway analysis.

Data file S2: Fibroblast cluster pathway analysis.

## Acknowledgments

We thank Implicit Bioscience Ltd (Brisbane, Australia) for supplying the murine anti-CD14 antibody (clone biG53) used in the DMM experiments, which was obtained under an MTA. We would like to thank Dr. Rui Xiao from the Department of Biostatistics and Epidemiology at the University of Pennsylvania for providing statistical guidance throughout study design and data analysis. We additionally thank the research staff at the CHOP CAG Sequencing Core, the Penn Center for Musculoskeletal Disorders (NIH/NIAMS P30AR069619) and the Chicago Center on Musculoskeletal Pain (C-COMP) (NIH/NIAMS P30AR079206) for their valuable contributions. We also acknowledge valuable contributions from Dr. Padmaja Mehta-D’souza, Taylor Conner, and Jessica Lumry from the Griffin Lab at OMRF.

## Author contributions

Conceptualization: KGB, SMS, TMG, AMM, REM, CRS.

Methodology: KGB, NSA, SI, SMS, VN, KLS, SYK, DBH, JFB, RLM, TMG, AMM, REM, CRS.

Investigation: KGB, NSA, SI, SMS, CZ, LYH, SYK, AER, BH, LAM.

Visualization: KGB, SMS, RLM, TMG, AMM, REM, CRS.

Funding acquisition: TMG, AMM, REM, CRS.

Project administration: KGB, BH, TMG, AMM, REM, CRS.

Supervision: KGB, RLM, TMG, AMM, REM, CRS.

Writing – original draft: KGB, RLM, CRS.

Writing – review & editing: KGB, SMS, NSA, VN, SI, CZ, KLS, SYK, AER, BH, LAM, LYHI, DBH, JFB, RLM, TMG, AMM, REM, CRS.

## Competing interests

AMM is consultant for Novartis, Merck, Roivant, and Averitas. She has received research support from Orion and Eli Lilly.

CRS and TMG are named inventors on US patent application pending entitled, “COMPOSITIONS AND METHODS FOR TREATING OSTEOARTHRITIS USING A CD14 INHIBITOR” (#18/411,242).

CRS is an Associate Editor for *Arthritis and Rheumatology*, and on the editorial boards of *Osteoarthritis & Cartilage*, and *OAC Open*.

TMG is consultant for Novo Nordisk.

## Data and materials availability

Datasets used for the present study are available from the corresponding author upon reasonable request.

