## Supplemental Materials for "CD14 plays a critical role in pain and inflammation across multiple models of post-traumatic osteoarthritis"

**Supplemental Materials and Methods**

*Synovial Fluid Samples:* Patients with knee OA who were eligible to receive knee injections and able to walk without assistive devices were recruited from orthopedic practices at four medical centers within the VA Health System across the continental United States (Omaha, Puget Sound, Washington D.C., and Philadelphia VA Medical Centers), under an Institutional Review Board (IRB) approved protocol. Full study criteria and methodology can be found at (*1*). Synovial fluid (SF) was then collected from patients receiving joint effusion prior to the intraarticular delivery of either lidocaine only, or lidocaine plus a corticosteroid. SF samples were received over a 6-month period, with multiple draws received per patient. Samples remained blinded for this study.

*Measurement of soluble CD14 in SF samples:*
SF was centrifuged at 500g to remove cell debris prior to storage at −80°C. Synovial fluid specimens were thawed, diluted 1: 500, and soluble CD14 was measured utilizing the Human CD14 Quantikine QuicKit ELISA (Cat#: QK383, R & D Systems, Minneapolis, MN).

*Measurement of knee hyperalgesia in clinical trial patients:* Knee hyperalgesia was measured in clinical trial patients (described above) receiving treatment for painful OA. Knee hyperalgesia was measured prior to initial joint effusion and therapeutic treatment, and again 6-wks following treatment. Pain pressure threshold values were reported as $kg/cm^2$, recorded from the average of three measurements taken at the knee joint using a handheld algometer.

*Animals:* Mice deficient in CD14 (B6.129S4-Cd14tm1Frm/J or CD14-/-, Jackson laboratory #003726) and wildtype (WT) C57BL/6 congenic controls (Jackson laboratory #000664) were obtained through Jackson laboratory. All animal procedures were approved by the Institutional Animal Care and Use Committees (IACUC) at the respective institutions where work was done (see below for each model). Model specific housing and procedures are described below.

*Destabilization of the medial meniscus (DMM) surgical model (mild PTOA):* All DMM model animals were housed at the Corp. Michael J. Crescenz VA Medical Center (CMCVAMC), and all experiments were approved by the Institutional Animal Care and Use Committees (IACUC). DMM mice were group-housed (4–5 animals/cage) in standard mouse cages held in a ventilated rack. Each cage was lined with woodchip bedding, animals were supplied with food and water ad libitum, and mice were provided with paper-mash nestlets. The room was kept on a 12:12 light:dark cycle. Destabilization of the medial meniscus (DMM) surgery was performed via resection of the medial meniscotibial ligament (MMTL) in skeletally mature (12-wk old) male mice to produce mild PTOA pathology, as previously described (*2, 3*). WT and CD14-KO mice that underwent DMM, and WT mice that underwent DMM and intra-articular delivery of IgG or anti-CD14 therapeutic, as described prior, were euthanized at 4- and 8-wks post injury. DMM mice were evaluated for spontaneous and evoked pain behaviors at 4-, and 8-wks, as detailed below. At euthanasia, DMM operated (right) and contralateral unoperated (left) knees were either isolated as whole joints for downstream evaluations in histopathology and imaging mass cytometry (IMC) or had anterior synovial and fat pad isolated for flow cytometry analysis, detailed below.

*Partial meniscectomy (PMX) surgical model (severe PTOA):* All PMX model animal procedures were approved by the Institutional Animal Care and Use Committees at Rush University Medical

Center and Northwestern University. PMX surgery was performed via removal of half of the medial meniscus in skeletally mature male and female mice to produce a severe PTOA pathology, as described (*4*). PMX mice were housed at Rush Medical Center with food (Teklad Global 18% Protein Rodent Diet, Inotiv #2018) and water ad libitum and kept on 12-h light cycles (light from 7am-7pm). PMX Mice were maintained in HEPA filtered ventilated micro-isolator racks using polycarbonate cages that have 75 in2 of floor space. PMX Mice were housed using Shepherd's® Cob PLUS™ (Shepherd Specialty Papers). Housing rooms were maintained at an average 45% humidity and 72 °F temperature. WT and CD14-KO mice underwent PMX and were euthanized at 10-wks post injury. PMX mice were evaluated for spontaneous and evoked pain behaviors at 4-, 6-, 8-, and 10-wks, as detailed below. At euthanasia PMX operated (right) and contralateral unoperated (left) knees were isolated as whole joints for downstream evaluations in histopathology, detailed below.

*High fat diet–anterior cruciate ligament rupture (HFD-ACLR) model (severe PTOA):* HFD-ACLR experiments were conducted following a protocol approved by the AAALAC-accredited Institutional Animal Care and Use Committees at Oklahoma Medical Research Foundation (OMRF) (protocol #20-30) and Oklahoma City VA (protocol #2001-001). Female C57BL/6N mice (Charles River #027) were housed at OMRF in specific pathogen-free conditions under a 14/10 light cycle and standard temperature conditions (21–23 °C) with ad libitum access to stock chow (5053, LabDiet) and chlorinated water (0.8–1.6 ppm), before being switched to a high fat diet (45% kcal fat, Research Diets D12451i) at 16-wks of age. Nonsurgical anterior cruciate ligament rupture (ACLR) was performed via externally applied compression overloading to a 1.7mm displacement (Bose ElectroForce 3100, Eden Prarie, MN, USA) in 36-wk old female HFD mice to produce a severe PTOA pathology, as described (*5*). ACLR was confirmed by auditory and mover-displacement cues. WT mice that underwent HFD-ACLR and intra-articular delivery of IgG or anti-CD14 therapeutic, as described prior, were euthanatized at 4-wks post injury. HFD-ACLR mice were evaluated for spontaneous and evoked pain behaviors at 2- and 4-wks, as detailed below. Following euthanasia, loaded (right) and contralateral unloaded (left) knees were isolated as whole joints for downstream evaluations in histopathology, detailed below.

*Drug intervention:* WT mice subjected to either DMM or ACLR-HFD were treated intra-articularly with either an anti-CD14 monoclonal antibody (mAb, clone biG53 LALA-PG, supplied by Implicit Bioscience in the DMM studies, or clone biG53 purchased from CellSciences, Cat# CMC005, in the ACLR-HFD studies) or an IgG2a control (LALA-PG-modified IgG2a supplied by Implicit Bioscience for use in the DMM studies, and clone B-Z2 purchased from CellSciences Cat#: CDM383A, for use in the ACLR-HFD studies), both at concentration of 0.5mg/kg. The LALA-PG modification reduces complement fixation and Fc-receptor binding, thus attenuating antibody-mediated cytotoxicity (*6*). For early treatment, mice received 3 weekly injections beginning 48 hours post DMM or ACLR injuries, described above. For the late delivery blockade cohort using the DMM model, WT mice began receiving 3-weekly injections beginning at 4-wks post-surgery.

*Knee hyperalgesia analysis:* Evoked Knee hyperalgesia was recorded using a pressure application measurement (PAM) device (Ugo Basile, Varese, Italy), measuring the hind paw withdrawal threshold in grams, as previously described (*7, 8*). For the DMM and PMX models, a maximum of 450 g of force was applied, and two measurements were recorded per knee and averaged, as described. If the mice did not withdraw their knee, the maximum force of 450 g was assigned. Knee hyperalgesia was measured at baseline and 4- and 8-weeks post DMM (n=6-12), at baseline and 2- and 4-weeks post ACLR injury (n=10), and at baseline and 4- and 8-weeks post PMX (n=5).

*Mechanical allodynia analysis:* PMX mice were tested for secondary mechanical allodynia of the ipsilateral hind paw using von Frey fibers and the up-down staircase method, as previously described (*8, 9*). The threshold force required to elicit withdrawal of the paw (median 50% withdrawal) was determined on each hind paw on each testing day. Withdrawal thresholds were measured at baseline and at 2- and 4-wks post PMX (n=5). For ACLR-HFD mice, a range of filaments were applied in ascending order, and the positive withdrawal response rate averaged across the first four filaments (as described in (*10*)). Withdrawal responses were measured at 2- and 4-wks post HFD-ACLR (n=10).

*Weight bearing analysis:* Paw weight bearing distribution was measured via the Advanced Dynamic Weight Bearing system (ADWB, Bioseb) or the static incapacitance meter system (Bioseb). For DMM and HDF-ACLR models ADWB was utilized to measure changes to paw weight bearing dynamics following injury, as previously described (*11*). For the DMM model, weight bearing dynamics were measured via the ADWB system at baseline and at 4- and 8-weeks post DMM surgery (n=6-12 mice per group). For the HFD-ACLR model, weight bearing was measured via the ADWB system at baseline and at 2- and 4-weeks post overloading (n= 10 mice per group).The static incapacitance meter system (Bioseb) was utilized to similarly measure changes to paw weight bearing dynamics following PMX injury, as previously described (*8*). Weight bearing asymmetry was measured via the static incapacitance meter system at baseline and 6- and 10-weeks post PMX surgery (n= 5 mice per group).

*Cage behavior analyses:* Spontaneous cage behavior was evaluated within the DMM model using the Laboratory Animal Behavior Observation Registration and Analysis System (LABORAS$^{TM}$, Metris) as previously described (*2*). Where cage activities such as climbing, rearing, and grooming were measured in addition to mobility metrics including total distance traveled within the cage, and average speed. Activity was measured at baseline (before DMM surgery) and at 4- and 8-weeks post DMM surgery (n=6-12 mice per group). Cage activity was also evaluated within the DMM and HFD-ACLR models using the Advanced Dynamic Weight Bearing system (ADWB, Bioseb), as previously described (*11*). Rearing activity measured via the ADWB system was measured at baseline and at 4- and 8-weeks post DMM (n=6-12 mice per group), and at baseline and 2- and 4-weeks post HFD-ACLR (n=10 mice per group).

*Histopathology:* At euthanasia, whole knee joints were isolated, fixed in 4% PFA (24 hrs), decalcified (14% EDTA at pH: 7.2, 7-days) and paraffin embedded. For DMM and PMX models evaluations of cartilage damage were performed on toluidine blue (proteoglycans and glycosaminoglycans) stained coronal sections taken midline through the joint using the modified Osteoarthritis Research Society International (OARSI) score as previously described (*12*). Average and max scores taken across medial and lateral tibial plateau and femoral condyle regions were reported. For DMM and PMX models features of synovial inflammation (synovitis) were evaluated, including lining hyperplasia, sublining cellularity, and fibrosis, using a murine specific scoring system on HE stained coronal sections taken midline through the joint (*13*). Feature specific scores were reported as the average across medial and lateral, femoral and tibial, synovial regions. For the HFD-ACLR model evaluations of cartilage damage were performed using the same scoring methods on hematoxylin, Safranin-O, and fast green stained sagittal sections (*12*). Average and max scores taken across femoral and tibial regions from medial and lateral compartments were reported. For the HFD-ACLR model the same synovitis scoring methods were performed hematoxylin, Safranin-O, and fast green stained sagittal sections (*13*). Feature specific scores were reported as the average across the anterior tibial synovium, posterior femoral synovium, and infrapatellar fatpad regions.

*Flow Cytometry:* Synovial and fat-pad tissue from 4 knees were pooled for each biological replicate (n=5 biological replicates/group/time point), collected at 0- (preop), 4-, 8- or 16-wks post-surgery, and cells were isolated enzymatically in Liberase (1U/mL) at 37°C for 30 minutes. Cell suspensions were split in half and stained for 30 minutes on ice with antibodies for monocyte (CD45-PerCP: Biolegend Cat #103130, Ly6C-APC: Biolegend Cat #128061), and macrophage ( CD64-PE: Biolegend Cat #139304) cell markers or T cell markers (CD45-PerCP, CD3-FITC: Biolegend Cat #100204, CD4-APC: Biolegend Cat #100412, CD8-PE: Biolegend Cat #100708). Cells were washed with FACs staining buffer (Biolegend Cat #420201) and stained with a fixable viability dye (Biolegend Cat #423114) for 30 minutes on ice. Cells were then washed again and resuspended in FACs staining buffer. Multicolor flow cytometry was performed (BD LSR II), and data was analyzed with FlowJo software (Version 10). Monocyte/macrophage populations were expressed as percent of the CD45+ population, T cell populations were expressed as percent of the CD45+ or CD3+ populations.

*Imaging Mass Cytometry (IMC):* Sagittal paraffin sections from the medial compartment of CD14 KO and WT, unoperated (contralateral) and DMM, knees 4-weeks post DMM surgery were utilized for IMC analysis, as previously described (n=4-6 mice per group) (*5*). Sections were deparaffinized and dehydrated in graded alcohol washes prior to heat mediated antigen retrieval within a citrate buffer (30 min at 99 °C) and overnight incubation at 4°C with a 24-panel metal-conjugated antibody cocktail (Supplemental Table 1). The following day slides were washed in PBS and stained with 125 μM Intercalator-Ir nuclear stain at room temperature for 30 min. Slides were then washed and air-dried for 20 min before loading into the Hyperion Imaging System (Standard BioTools, South San Francisco, CA, USA). Regions of interest (ROI) spanning 350 μm x 350μm and containing posterior- and anterior- femoral and tibial synovial gutters were hand selected for laser ablation based on transmitted light images. Raw mass cytometry data was acquired for each 1 $\mu m^2$ pixel within ROIs to generate MCD image files. MCD files were visualized using MCD Viewer (Standard Biotools) and exported from the MCD viewer as a multi-OME tiff of all working metal conjugated antibody channels and the 125μM Intercalator-Ir nuclear stain, DNA, channel. Single cell masks were created the DNA channel only tiffs (deepcell.org). To minimize metal contamination for all solutions were prepared using mass spectrometry grade Maxpar (Standard Biotools) water and PBS, optimized for the best signal-to-noise ratio.

*Spatial protein expression and cellular phenotype analysis:* Cell masks and Multi-OME tiff files for each individual ROI were loaded into IMACytE software, to create t-distributed stochastic neighbor embedding (t-SNE) dimensionality reduction analysis with arcsine transformation to produce data normalization and cluster analysis, as previously described (*5, 14*). Cluster marker expression heatmaps, cell cluster assignment ROI images, and cell counts per cluster within each ROI was exported. Neighborhood analysis was performed within IMACyte, identifying nearest neighboring clusters found within 10µm of each other, with >5 interactions considered significant. Neighbhorhood analysis was reported via glpyhs of neighbhoring cell interactions and summary statistics (Z-score), as previously described (*5*).

*Single-cell RNA-sequencing (scRNA-seq):* For scRNA-seq synovial tissue was isolated and pooled from the right knees of 15 male WT and CD14 KO mice that had undergone DMM surgery 4-wks prior. Synovial tissue was digested in RPMI containing Liberase (1 µU/mL) and DNase I (200µg/mL), for 30 min at 37°C on a shaker, and vortexed every 10 minutes. Tissue digestion was stopped by the addition of 10% fetal bovine serum FBS. Following digestion, synovial single cell suspensions were split into two for a technical duplicate, and immediately submitted to the Children's Hospital of Philadelphia (CHOP) Center for Applied Genomics (CAG) Sequencing Core for initial quality check to ensure cell suspension viability > 90%. Next-generation sequencing libraries were prepared using the 10x Genomics Chromium Single Cell 3' Reagent kit v3.1 according to the manufacturer's instructions. Libraries were uniquely indexed using the Chromium Dual Index Kit, pooled, and sequenced on an Illumina NovaSeq 6000 sequencer in a paired-end, dual indexing run. Sequencing for each library targeted 20,000 mean reads per cell. Data were then processed using the Cell Ranger pipeline (10x Genomics, v.6.1.2) for demultiplexing, alignment of sequencing reads to the mm10 transcriptome, and creation of feature-barcode matrices.

*scRNA-seq data quality control and analysis*

Aligned data from Cell Ranger was then read from Cell Ranger into Seurat (R, v4.1.0). Low quality cells were filtered out based on the number of molecules, with cells < 200 molecules and in the bottom 8% based off molecule count. Further, cells were filtered out based on the number of genes (Seurat "features"), with cells < 350 features and in the bottom 8% based off molecule count. Cells with > 5% of all genes being mitochondrial-derived were also excluded. Following quality control, raw gene counts were normalized, and data was merged across experimental groups. Uniform Manifold Approximation and Projection (UMAP) dimensions 1-20 were used in the object representing all groups and cells. Unsupervised clustering was performed using Seurat *FindNeighbhors* with dimensions 1-20 and Seaurat *FindClusters* with a resolution of 0.16 to identify distinct cell clusters. Clusters were identified using Seurat *FindAllMarkers*, reporting genes expressed in greater than 90% of cells in the cluster of interest and less than 10% of cells in all other clusters. Only genes with a positive differential expression relative to other clusters were included. To further identify cell type subsets (i.e. monocytes) all monocyte-like clusters were subset from all sequenced cells and re-clustered (dimensions 1-20, resolution 0.16). Again, *FindAllMarkers* was applied within the re-clustered groups as described above and gene lists were uploaded to Reactome pathway analysis to compare gene marker profiles with known cellular pathways (*15*). Total cell numbers per cluster within groups were plotted.

**REFFERENCES:**

1. Leach, W., C. Doherty, M. Olave, B.R. England, K. Wysham, G. Kerr, M. Quinones, A. Ogdie, D. White, and T. Neogi, *Protocol for a multi-center randomized controlled trial to evaluate the benefits of exercise incentives and corticosteroid injections in osteoarthritis of the knee (MOVE-OK).* Trials, 2022. **23**(1): p. 604.

2. Sambamurthy, N., C. Zhou, V. Nguyen, R. Smalley, K.D. Hankenson, G.R. Dodge, and C.R. Scanzello, *Deficiency of the pattern-recognition receptor CD14 protects against joint pathology and functional decline in a murine model of osteoarthritis.* PLoS One, 2018. **13**(11): p. e0206217.
3. Glasson, S., T. Blanchet, and E. Morris, *The surgical destabilization of the medial meniscus (DMM) model of osteoarthritis in the 129/SvEv mouse.* Osteoarthritis and cartilage, 2007. **15**(9): p. 1061-1069.
4. Knights, C.B., C. Gentry, and S. Bevan, *Partial medial meniscectomy produces osteoarthritis pain-related behaviour in female C57BL/6 mice.* Pain, 2012. **153**(2): p. 281-292.
5. South, S.M., M.C. Marlin, P. Mehta-D'souza, T. Stephens, T. Conner, K.G. Burt, J.M. Guthridge, C.R. Scanzello, and T.M. Griffin, *Imaging mass cytometry reveals tissue-specific cellular immune phenotypes in the mouse knee following ACL injury.* Osteoarthritis and Cartilage Open, 2023. **5**(4): p. 100416.
6. Lo, M., H.S. Kim, R.K. Tong, T.W. Bainbridge, J.-M. Vernes, Y. Zhang, Y.L. Lin, S. Chung, M.S. Dennis, and Y.J.Y. Zuchero, *Effector-attenuating substitutions that maintain antibody stability and reduce toxicity in mice.* Journal of Biological Chemistry, 2017. **292**(9): p. 3900-3908.
7. Miller, R.E., S. Ishihara, B. Bhattacharyya, A. Delaney, D.M. Menichella, R.J. Miller, and A.M. Malfait, *Chemogenetic inhibition of pain neurons in a mouse model of osteoarthritis.* Arthritis & Rheumatology, 2017. **69**(7): p. 1429-1439.
8. Obeidat, A.M., M.J. Wood, N.S. Adamczyk, S. Ishihara, J. Li, L. Wang, D. Ren, D.A. Bennett, R.J. Miller, and A.-M. Malfait, *Piezo2 expressing nociceptors mediate mechanical sensitization in experimental osteoarthritis.* Nature communications, 2023. **14**(1): p. 2479.
9. Miller, R.E., P.B. Tran, R. Das, N. Ghoreishi-Haack, D. Ren, R.J. Miller, and A.-M. Malfait, *CCR2 chemokine receptor signaling mediates pain in experimental osteoarthritis.* Proceedings of the National Academy of Sciences, 2012. **109**(50): p. 20602-20607.
10. Komaravolu, R.K., P. Mehta-D'souza, T. Conner, M. Allen, J. Lumry, A. Batushansky, N.P. Pezant, C.G. Montgomery, and T.M. Griffin, *Sex-specific effects of injury and beta-adrenergic activation on metabolic and inflammatory mediators in a murine model of post-traumatic osteoarthritis.* Osteoarthritis and Cartilage, 2024. **32**(9): p. 1097-1112.
11. Dent, J.O., J.P. Segal, A. Brécier, H.G. Gowdy, R.M. Dubois, C.A. Bannerman, K. Halievski, J.R. Silva, and N. Ghasemlou, *Advanced dynamic weight bearing as an observer-independent measure of hyperacute hypersensitivity in mice.* Canadian Journal of Pain, 2023. **7**(2): p. 2249060.
12. Glasson, S., M. Chambers, W. Van Den Berg, and C. Little, *The OARSI histopathology initiative–recommendations for histological assessments of osteoarthritis in the mouse.* Osteoarthritis and cartilage, 2010. **18**: p. S17-S23.
13. Obeidat, A.M., S.Y. Kim, K.G. Burt, B. Hu, J. Li, S. Ishihara, R. Xiao, R.E. Miller, C. Little, and A.-M. Malfait, *A Standardized Approach To Evaluation And Reporting of Synovial Histopathology In Two Surgically Induced Murine Models Of Osteoarthritis.* Osteoarthritis and Cartilage, 2024.
14. Somarakis, A., V. Van Unen, F. Koning, B. Lelieveldt, and T. Höllt, *ImaCytE: visual exploration of cellular micro-environments for imaging mass cytometry data.* IEEE transactions on visualization and computer graphics, 2019. **27**(1): p. 98-110.
15. Milacic, M., D. Beavers, P. Conley, C. Gong, M. Gillespie, J. Griss, R. Haw, B. Jassal, L. Matthews, and B. May, *The reactome pathway knowledgebase 2024.* Nucleic acids research, 2024. **52**(D1): p. D672-D678.

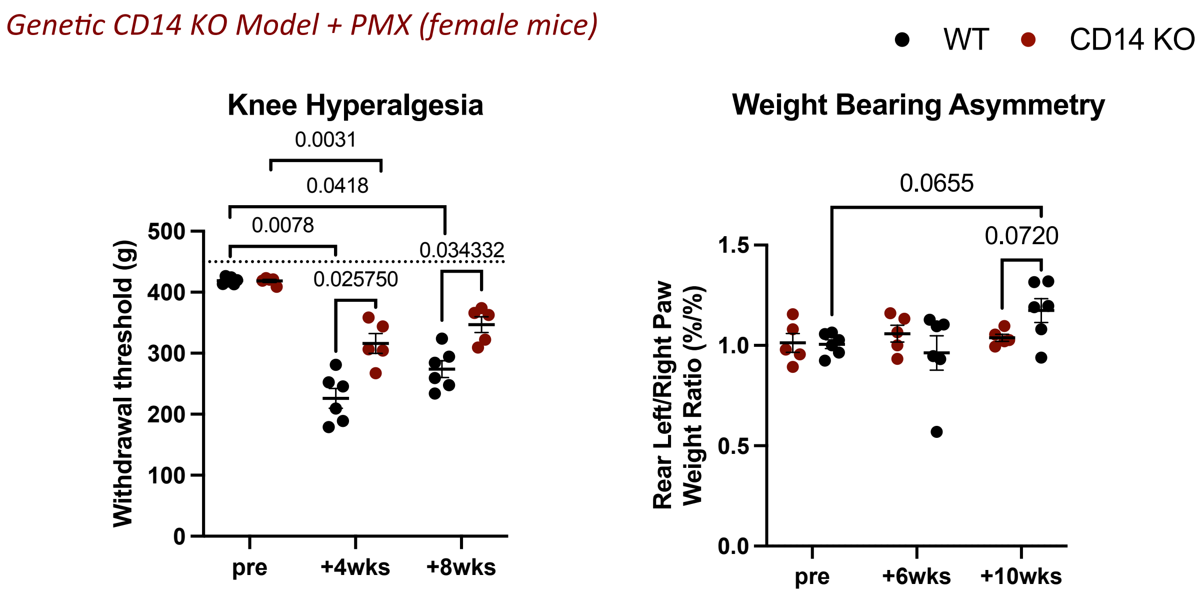


**Fig. S1: Pain and weight bearing changes between WT and CD14 KO female mice following PMX injury.** Knee hyperalgesia measured in WT (n=6) and CD14 KO (n=5) mice via withdrawal threshold (g) using a PAM device at pre-PMX and 4- and 8-wks post PMX. Weight bearing asymmetry measured via rear right to rear left paw weight ratio (%/%) using the static incapacitance meter system at pre-PMX and 6- and 10-wks post PMX. Statistically significant differences in knee hyperalgesia were calculated using the Mann-Whitney U non-parametric T-test with Holm-Šídák post hoc, and across time points using the Friedman test with Dunn's multiple comparison adjustment. Statistically significant differences in weight bearing asymmetry were calculated using a repeated measures two-way ANOVA followed by Šídák post-hoc for multiple comparisons.

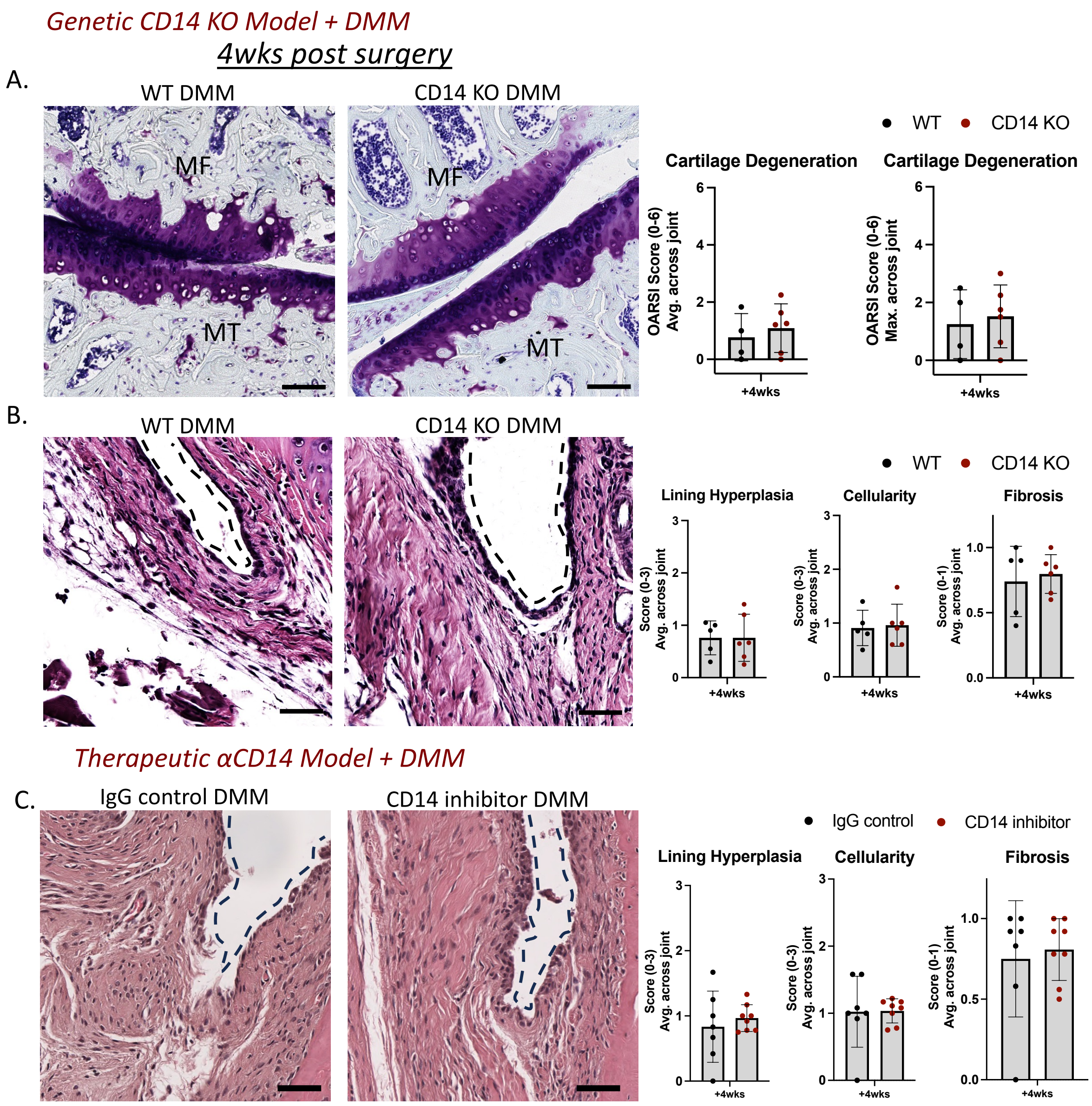


**Fig. S2: Histopathologic evaluations of cartilage damage and synovitis at intermediate time points withing CD14 KO and CD14 blockade DMM models.** (**A**) Representative images of Toluidine Blue stained histological sections of WT (n=4) and CD14 KO (n=6) tibial and femoral cartilage surfaces 4-wks following DMM (MF: medial femoral, MT: medial tibial) (Scale bar = 100μm). Associated cartilage scoring measured as the average or max scores reported across the medial and lateral, femoral and tibial cartilage surfaces. (**B**) H&E stained histological sections of WT (n=5) and CD14 KO (n=6) medial femoral synovial gutters 4-wks following DMM (Scale bar = 50μm). Associated synovitis scoring measured as the average scores reported across synovial regions within the joint. (**C**) H&E stained histological sections between IgG (n=7) or CD14 inhibitor (n=8) treated WT medial femoral synovial gutters 4-wks following DMM (Black dashed line indicates synovium lining) (Scale bar = 50μm). Associated synovitis scoring measured as the average scores reported across synovial regions within the joint. Statistically significant differences were calculated using a student's T-test.

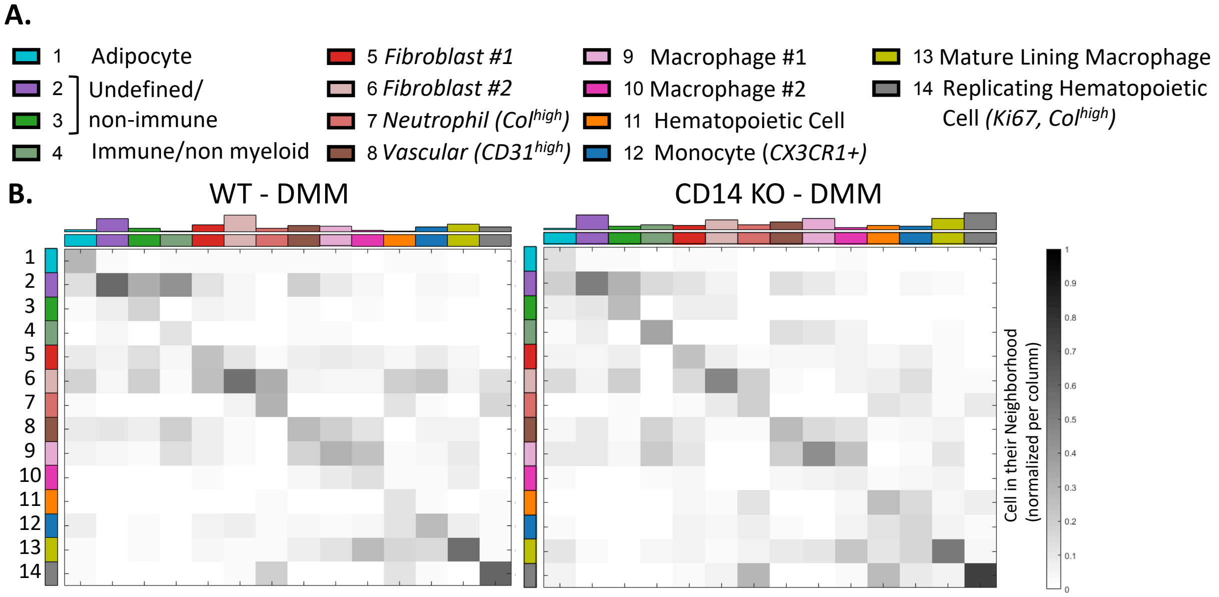


**Fig. S3: Neighborhood analysis of IMC cell clusters across WT and CD14 KO synovium 4-wks post DMM.** (**A**) IMC cell clusters. (**B**) Cell cluster interaction heatmap normalized per column. The relative frequency (grayscale) that a cell cluster of interest (column) is associated with another specific cell cluster micro-environment (row). Bar chart along top of heatmap represents the total number of cells across ROIs within each cluster of interest.

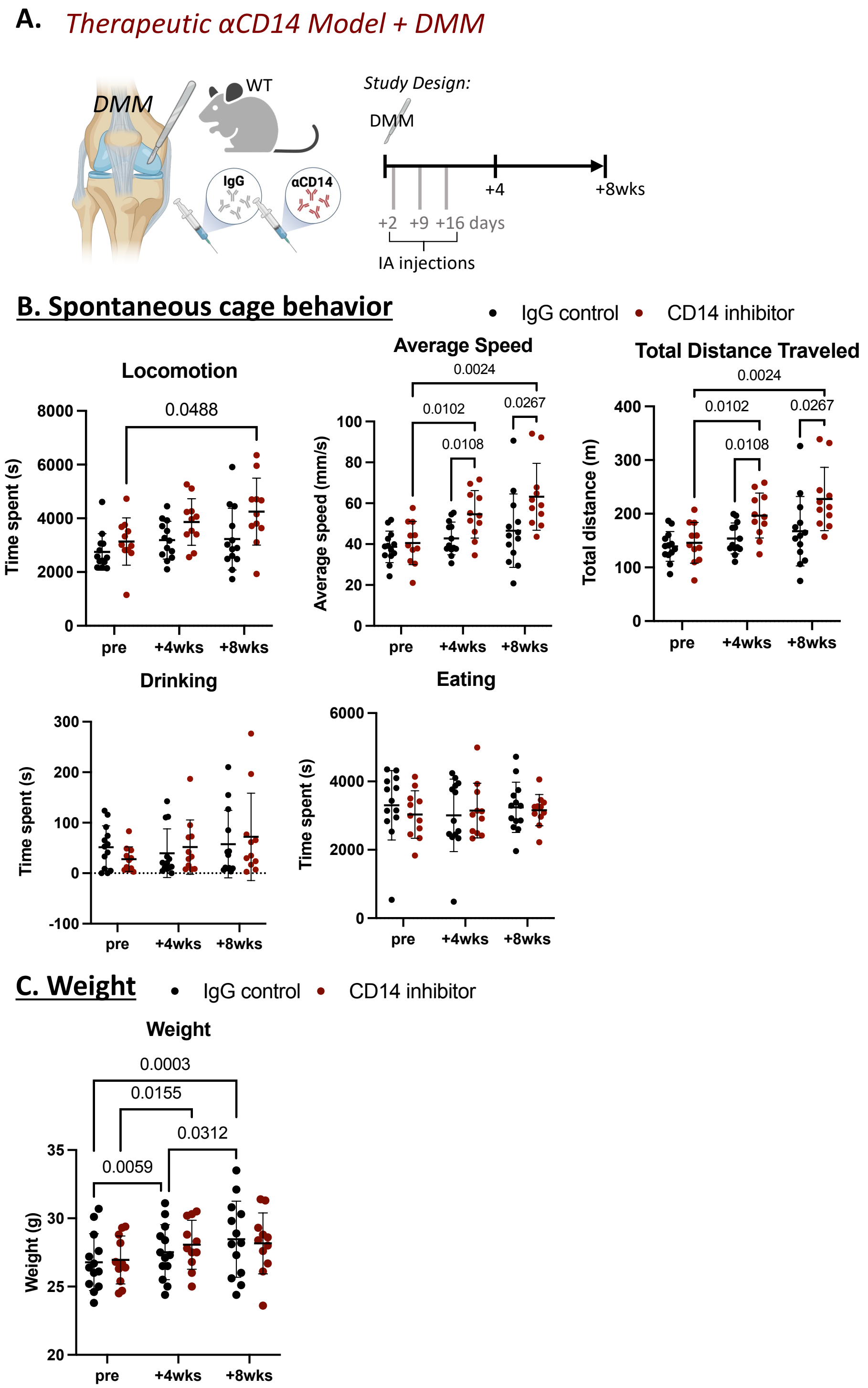


**Fig. S4: Behavioral data from early CD14 blockade delivery DMM model.** (**A**) Study schematic of the early delivery of IgG (n=13) or CD14 inhibitor (n=11) treated WT mice following DMM surgery. (**B**) Cage activity measured as the time spent (s) in locomotion, average speed of mise (mm/s), total distance traveled (m), time spent drinking (s), and time spent eating (s) using the ADWB system. (**C**) Animal weights (g) throughout study. Statistically significant differences were calculated using a use repeated measures mixed effect model followed by Šídák post-hoc for multiple comparisons.

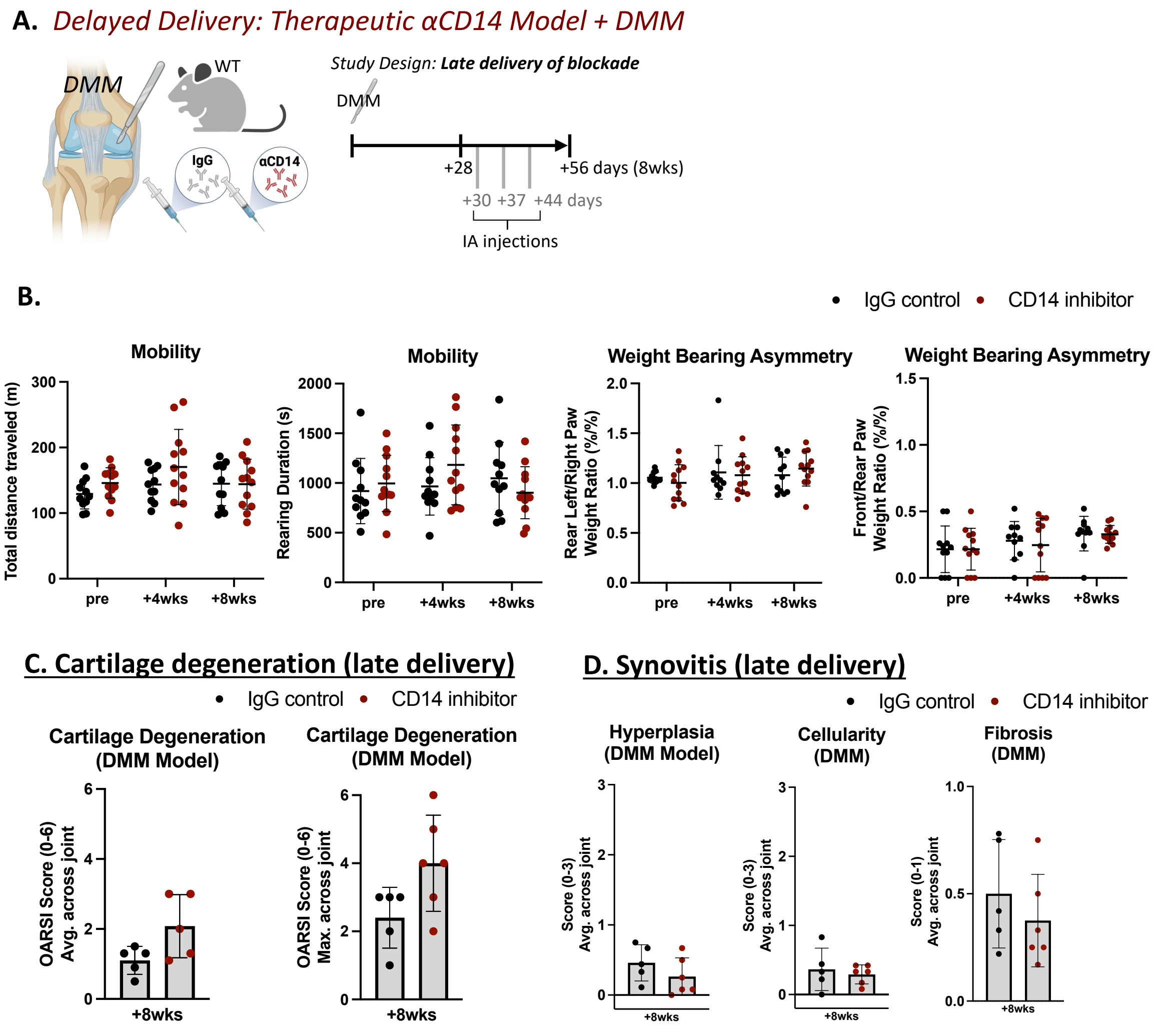


**Fig. S5: Pain, behavioral, and histopathologic evaluations within the late CD14 blockade delivery DMM model.** (**A**) Study schematic of the late delivery of IgG (n=11) or CD14 inhibitor (n=12) treated WT mice following DMM surgery. (**B**) Mobility measured via total distance traveled (m) and rearing duration (s) using the ADWB system. Weight bearing asymmetry measured as the front to rear paw weight ratio (%/%) and rear-left to rear-right paw weight ratio (%/%). (**C**) Cartilage scoring of IgG (n=5) or CD14 inhibitor (n=6) femoral and tibial cartilage 8-wks post DMM surgery, measured as the average or max scores reported across the medial and lateral, femoral and tibial cartilage surfaces. (**D**) Synovitis scoring of synovial tissue 8-wks post DMM surgery, measured as the average scores reported across synovial regions within the joint. Statistically significant differences in behavior outcomes (**B**) were calculated using a repeated measures mixed effect model followed by Šídák post-hoc for multiple comparisons. Statistically significant differences in histopathological scoring (**C,D**) were calculated using a student's T-test.

**Table S1: Synovial fluid patient characteristic information.** Information on the number of synovial fluid draws, sex, age, and body mass index (BMI) of patients from which synovial fluid and pain behavior was analyzed.

| CD14 Study Patient ID | Draws | Sex | Age | BMI |
|---|---|---|---|---|
| 1 | 2 | male | 64 | 29.4 |
| 2 | 2 | male | 79 | 31.9 |
| 3 | 2 | male | 73 | 33.4 |
| 4 | 2 | male | 51 | 35.9 |
| 5 | 2 | male | 76 | 25.2 |
| 6 | 2 | male | 67 | 30.1 |
| 7 | 2 | male | 53 | 39.1 |
| 8 | 2 | male | 68 | 23.9 |
| 9 | 2 | male | 60 | 45.4 |
| 10 | 2 | male | 74 | 30.1 |
| 11 | 2 | female | 65 | 33.1 |
| 12 | 2 | male | 72 | 36 |
| 13 | 2 | male | 69 | 29.2 |
| 14 | 2 | male | 70 | 28.5 |

**Table S2: List of antibodies used for IMC.** Pre-conjugated metal tagged antibodies acquired from Standard Biotools, additional custom conjugated antibodies were generated using commercially available antibodies with Standard Biotools custom conjugation kit. Concentration, vendor, and catalog #'s are listed.

| Metal | Target | Concentration (mg/mL) | Vendor | Catalog# |
|---|---|---|---|---|
| 141Pr | aSMA | 0.005 | Standard Biotools | 3141017D |
| 143Nd | Vim | 0.005 | Standard Biotools | 3143027D |
| 144Nd | CD14 | 0.005 | Standard Biotools | 3144025D |
| 168Er | Ki-67 | 0.005 | Standard Biotools | 3168022D |
| 169Tm | Col1a | 0.0025 | Standard Biotools | 3169023D |
| 170Er | CD3 | 0.005 | Standard Biotools | 3170019D |
| 162Dy | CD4 | 0.005 | Standard Biotools | 91H031162 |
| 176Yb | CD8 | 0.005 | Standard Biotools | 91H023176 |
| 171Yb | CD31 | 0.005 | Standard Biotools | 91H027171 |
| 150Nd | Ly6G | 0.001 | Standard Biotools | 91H037150 |
| 151Eu | CD45 | 0.005 | Standard Biotools | 91H029151 |
| 156Gd | F4/80 | 0.005 | Standard Biotools | 91H030156 |
| 158Gd | MHC-II (I-A/I-E) | 0.005 | Standard Biotools | 91H038158 |
| 142Nd | Ly6C | 0.005 | Biolegend | 128039 |
| 148Nd | CX3CR1 | 0.01 | Biolegend | 149002 |
| 149Sm | Cadherin 11 | 0.005 | Genetex | GTX635978 |
| 160Gd | Tenascin C | 0.005 | Novus Biologics | NB110-68136 |
| 154Sm | Perilipin | 0.005 | Abcam | ab3526 |
| 153Eu | CD56 | 0.005 | GeneTex | GTX634792 |
| 164Dy | CD24 | 0.005 | BIolegend | 101829 |
| 165Ho | Mast Cell Tryptase | 0.001 | Genetex | GTX75011 |
| 166Er | CD64 | 0.0025 | Biolegend | 161002 |
| 173Yb | Col2a1 | 0.005 | RnD | AF3615 |
| 175Lu | PGP9.5 | 0.005 | Abcam | ab27053 |
